# Spectral decompositions of neural voltage recordings are susceptible to model misspecifications that cause meaningful estimation error

**DOI:** 10.64898/2026.06.12.718232

**Authors:** Patrick F. Bloniasz, Emily P. Stephen

**Affiliations:** Graduate Program for Neuroscience, Boston University, Boston, MA 02215, USA; Department of Mathematics and Statistics, Boston University, Boston, MA 02215, USA

**Keywords:** power spectra, spectral decomposition, aperiodic activity, neural field potentials, propofol anesthesia, model misspecification

## Abstract

The power spectra of neural voltage recordings vary systematically across brain states and contain both narrowband (rhythmic) and broadband components. A large class of algorithms seeks to parametrize these spectra by separating rhythms from broadband structure, enabling many robust empirical findings. Here we show that two common assumptions underlying popular spectral decomposition methods are incompatible with standard physical and statistical properties of neural recordings: (1) field potentials arise from additive (linear) superposition of biophysical processes, yet several methods implicitly impose multiplicative structure; (2) power estimates are Gamma distributed, with variance proportional to squared power (heteroscedasticity), yet many methods assume Gaussian, homoscedastic errors across frequencies. Using simulations with known ground truth, we demonstrate how these misspecifications bias estimates of rhythm amplitude and broadband height/slope, even under well-behaved conditions. We introduce a corrected decomposition framework, released as the open-source package `SL_specdecomp`. Relative to the most widely used method, `specparam`, our approach recovers rhythms and broadband parameters accurately, while `specparam` decompositions are biased and can confound rhythmic peaks with broadband slope. We then apply these methods to monkey electrocorticography during propofol anesthesia. `SL_specdecomp` estimates a substantially steeper (more negative) 40–60 Hz broadband slope during anesthesia than during wakefulness, whereas `specparam` shows a smaller state difference. We show using simulation that the differences in the two decompositions can arise directly from `specparam`’s model misspecification. We also introduce a formal method based on cross-validated log likelihood to compare candidate power spectral decompositions and show that it favors `SL_specdecomp`. These results suggest that misspecified decompositions can attenuate or distort broadband slope changes in the presence of strong rhythms, and motivate the use of `SL_specdecomp` as a more reliable decomposition tool.

## 1 Introduction

Neural electrophysiological voltage recordings (e.g., electrocorticography, ECoG; electroencephalography, EEG) have been shown to contain rhythmic and broadband information (Donoghue, 2025; Donoghue & Watrous, 2023; Manning et al., 2009) that separately provide reliable biomarkers of healthy and disordered brain states (Donoghue, 2025; Donoghue & Watrous, 2023; Donoghue et al., 2020; Gerster et al., 2022). Rhythms can be transient or sustained (Jones, 2016; Muthuswamy & Thakor, 1998), and are typically attributed to resonance in the underlying network dynamics, leading to synchronous rhythmic population activity. Broadband power is thought to arise from asynchronous or stochastic population activity (Bloniasz et al., 2025; Brake et al., 2024; Gao, 2016; Gao et al., 2017; Halnes et al., 2024; Miller et al., 2009; Perrenoud & Cardin, 2023). Thus, broadband effects are often interpreted in terms of changes in average firing rate (e.g., Gao, 2016; Gao et al., 2017; Manning et al., 2009) or changing relative contributions of processes with different intrinsic time scales (Gao et al., 2020; Lendner et al., 2024). Standard power spectral estimates, which are the decomposition of a signal’s variance as a function of frequency (Kass et al., 2014; D. B. Percival & Walden, 1993; Priestley, 1982), superimpose rhythmic and broadband information, so changes in one can appear as changes in the other, complicating physiological interpretation.

Over the past decade, a growing class of algorithms has sought to separate these components parametrically—what we term power spectral decomposition methods (Beck et al., 2023; Donoghue et al., 2020; Hu et al., 2024; Matsuda & Komaki, 2017; Medrano et al., 2025; Wen & Liu, 2016). The most widely used of these, `specparam` (formerly “FOOOF”; Donoghue et al., 2020), has been cited in over two thousand studies since its 2020 release. Yet despite the ubiquity of these algorithms, the physical and statistical assumptions underlying spectral decomposition remain largely understudied. Here, we consider spectral decomposition as an estimation problem, analyzing its behavior using statistically grounded neural simulations and well-characterized ECoG data.

We identify and study two systematic forms of model misspecification that affect many current methods, including `specparam`. First, while neural field potentials are generally modeled as additive superpositions of biophysical processes such as synaptic and action potentials (Bloniasz et al., 2025; Buzsáki et al., 2012; Destexhe & Bédard, 2022; Nunez & Srinivasan, 2006), several decomposition algorithms implicitly assume multiplicative interactions (Brady & Bardouille, 2022; Donoghue et al., 2020). Second, because empirical power spectra derived from Gaussian time-domain signals follow Chi-square or Gamma distributions (D. B. Percival & Walden, 1993), their variance scales with frequency (heteroscedasticity). Nonetheless, most methods assume Gaussian, homoscedastic probability models (Donoghue et al., 2020; Medrano et al., 2025). Here we study the consequences of these model misspecifications and propose a corrected framework for spectral decomposition.

First, we demonstrate in simulation that these misspecifications can yield erroneous rhythmic–broadband decompositions even under well-behaved conditions, using `specparam` as a case study. Then, we argue that these same errors occur in real data, using electrocorticography recordings in monkeys during propofol anesthesia. In the process, we provide 1) a simulation tool with known ground truth to test existing or future algorithms (SL_GPsim), 2) a method using cross-validated log likelihood to compare candidate spectral decompositions, and 3) a decomposition software package (SL_specdecomp) where we use a corrected probability model for the power spectra and give the option to use either multiplicative or additive superposition of components. Together, our results call for cautious reinterpretation of prior findings derived from misspecified models and motivate the use of our tools for spectral decomposition, model validation, and model comparison.

## 2 Background

### 2.1 Additive generative structure of neural field potentials

Under the standard model of local field potentials (LFPs), the extracellular voltage recorded at a point in the brain arises from the linear superposition of transmembrane currents generated by nearby neurons. Each current source contributes additively to the potential through the resistive extracellular medium. Because the extracellular medium behaves approximately as a homogeneous resistor at frequencies below ∼1 kHz, this superposition is linear and instantaneous—a property preserved across cortical and subcortical structures. Thus, the observed potential *V* (*t*) is the sum of many elementary current generators (synaptic, dendritic, and spiking), each filtered by biophysical and geometric factors (briefly reviewed in Bloniasz et al., 2025, see also Bédard et al., 2004; Buzsáki et al., 2012; Halnes et al., 2024; Nunez and Srinivasan, 2006; Rall and Shepherd, 1968): 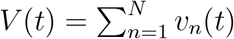.

In the frequency domain, this linearity implies that the total power spectrum of the field potential is linear. For example, if the current generators are independent, the power spectrum is a sum:

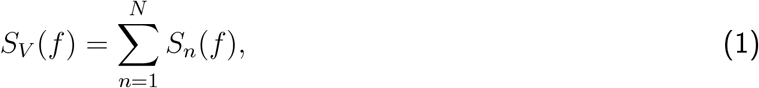

where *S*_*n*_(*f* ) is the power spectral density (power spectrum) of the *n*-th current generator.

More generally, if the generators are dependent then there will be additional cross-terms in the summation:

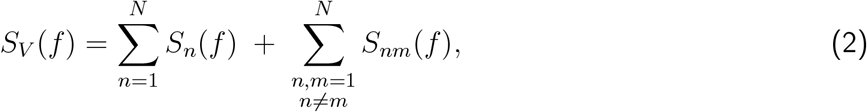

where *S*_*nm*_(*f* ) are cross-spectra capturing coherence between generators (Bloniasz et al., 2025).

Importantly, the terms *broadband* and *narrowband* refer to spectral phenomenology (1/*f* -like backgrounds versus localized peaks), not to unique biophysical classes of input. Synaptic and spiking currents can each contribute power across wide frequency ranges, and structured network dynamics can generate narrowband peaks. One popular theoretical modeling approach, filtered point processes or shot noise, proposes that neural events like synaptic potentials and action potentials contribute both narrowband and broadband components: broadband components come from the stochastic timing of the events, which behave like filtered white noise, while narrowband components come from mean-field dynamics of the population event rates (Bloniasz et al., 2025; Brake et al., 2024; Destexhe & Bédard, 2022; Gao, 2016; Gao et al., 2017; Halnes et al., 2024; Miller et al., 2009; Perrenoud & Cardin, 2023). Under this framework, despite the fact that the same event process (synapses, action potentials) generates both narrowband and broadband components, the narrowband and broadband power spectral components sum together as in Eq. (1)(Bédard et al., 2006; Bloniasz et al., 2025; Destexhe & Bédard, 2022; Halnes et al., 2024).

In general, spectral decomposition methods aim to decompose *S*_*V*_ (*f* ) into components with distinct spectral shapes (broadband background and oscillatory peaks) while often remaining agnostic about the precise cellular origin of each component. A common decomposition writes the spectrum as the sum of a broadband term and one or more oscillatory (rhythmic) terms,

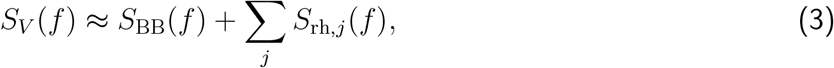

which corresponds to assuming that the broadband generator(s) and rhythmic generator(s) are approximately uncorrelated so that their cross-spectra are negligible. Here we adopt this independence/zero-coherence approximation for simplicity; coherent generators can be incorporated by retaining the appropriate interaction (cross-spectral) terms in Eq. (2) (Bloniasz et al., 2025; Gao et al., 2017; Halnes et al., 2024; Milstein et al., 2009).

This additive framework motivates spectral decomposition methods that treat broadband and rhythmic contributions as separable sources of power. In contrast, multiplicative models of the power spectrum, which combine components in log-power space (e.g., *S*_*V*_ (*f* ) = *S*_BB_(*f* ) × *S*_rh_(*f* )), implicitly assume that one process scales the amplitude of another. Such multiplicative interactions contradict the typically assumed linearity of field potentials and introduce model misspecification under this assumption.

### 2.2 Power spectra of Gaussian processes are Gamma distributed

Neural field potentials are often well approximated by Gaussian processes, particularly during stable resting states (e.g., (Elul, 1969)). Models of field potentials typically adopt this assumption, either by introducing additive Gaussian process noise (e.g., Borgers, 2017) or by employing biophysical formulations whose limiting dynamics converge to Gaussian processes, such as filtered point processes (e.g., Bloniasz et al., 2025; Gao et al., 2017; Halnes et al., 2024; Milstein et al., 2009).

A Gaussian process is fully specified by its mean and autocovariance, both of which may change in time. In practice, field potentials are typically analyzed under a local stationarity assumption within a chosen time window, so that the mean is constant and the autocovariance depends only on lag: 𝔼[*x*(*t*)] = *µ* and *C*_*x*_(*τ* ) = Cov(*x*(*t*), *x*(*t* + *τ* )). For a mean-zero, wide-sense stationary process, the true power spectrum *S*_*x*_(*f* ) is defined as the Fourier transform of the autocovariance (Wiener–Khinchin theorem),

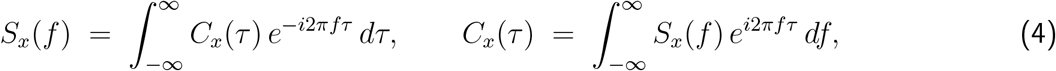

so that *S*_*x*_(*f* ) can be interpreted as a decomposition of the signal variance across frequencies (indeed, *C*_*x*_(0) = Var[*x*(*t*)] = ∫ *S*_*x*_(*f* ) *df* under standard conventions). Thus, a stationary Gaussian process may be equivalently characterized by (*µ, C*_*x*_(*τ* )) or by (*µ, S*_*x*_(*f* )). The goal in spectral estimation is to estimate *S*_*x*_(*f* ) from a finite sample.

As a concrete example, the Ornstein–Uhlenbeck (OU) process is the continuous-time analogue of a first-order autoregressive (AR(1)) Gaussian process, defined by:

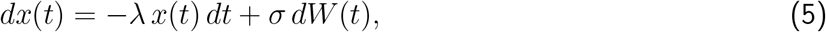

where *dW* (*t*)*/dt* can be thought of as white noise and *λ* > 0. When stationary, an OU process has autocovariance 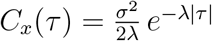 and corresponding “Lorentzian” spectrum:

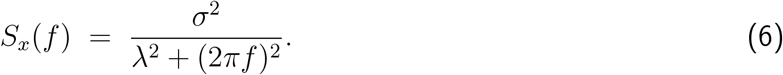

Hence a stationary OU process is a Gaussian process with zero mean and the spectrum in Eq. 6. These two elements completely describe the process in a stationary state.

Here we will show that when estimating the power spectrum of a Gaussian process using direct spectral estimators (such as the periodogram, Welch’s method, or multitaper spectral estimation), the spectral estimates will be Gamma distributed. Let 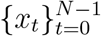 be a mean-zero, stationary Gaussian process sampled at interval Δ*t* (sampling rate 1*/*Δ*t*), and let 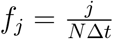 denote the Fourier frequencies, *j* = 1, …, ⌊*N/*2⌋. Following standard normalizations for direct spectral estimators (Kass et al., 2014; Kramer & Eden, 2016; D. B. Percival & Walden, 1993), define a tapered discrete Fourier transform (DFT):

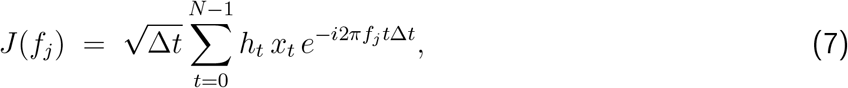

where {*h*_*t*_} is a data taper (with the untapered case recovered by *h*_*t*_ ≡ *N* ^−1/2^). Under standard large-sample conditions (large *N* so that there is negligible spectral leakage and distinct Fourier coefficients are approximately uncorrelated), the tapered Fourier coefficients are circularly symmetric complex normal with variance equal to the true spectrum,

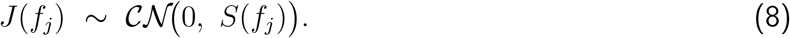

A direct estimate for *S*(*f*_*j*_ ) is the magnitude square of the Fourier coefficient, |*J* (*f*_*j*_ )|^2^. Many practical estimators reduce variance by averaging across *K* uncorrelated direct estimates; e.g., across orthogonal Slepian tapers in multitaper estimation, across segments in Welch’s method, or across independent trials:

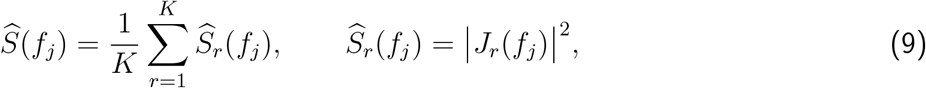

where *r* indexes the uncorrelated direct estimates being averaged. A classical result (Kass et al., 2014; Kramer & Eden, 2016; D. B. Percival & Walden, 1993) is that these averaged direct spectral estimators have a chi-square sampling distribution with 2*K* degrees of freedom:

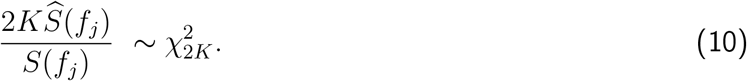

This result comes directly from the fact that the chi-square distribution captures sums of squared independent standard normal random variables; the factor of 2 comes from the two degrees of freedom in each complex normal Fourier coefficient.

The chi-square distribution is a special case of the Gamma distribution, where:

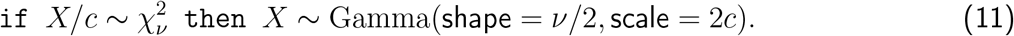

Setting 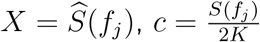, and *ν* = 2*K*, Eq. (10) becomes:

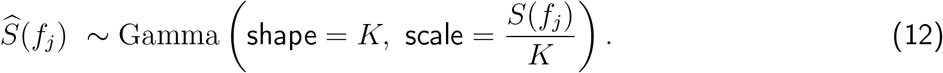

Hence, averaged direct spectral estimates of Gaussian processes are Gamma distributed. Here *K* denotes the number of uncorrelated direct estimates being averaged. For a multitaper spectrum averaged only across independent tapers, *K* = *n*_tapers_; for Welch’s method, *K* is the number of uncorrelated segments; for trial-averaged spectra, *K* is the number of independent trial spectra being averaged. More generally, *K* can be interpreted as the effective number of uncorrelated direct estimates contributing to the average.

One important consequence of this sampling distribution is that:

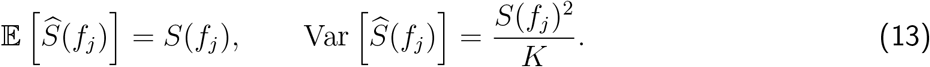

In other words, when *N* is large, the spectral estimates are unbiased but intrinsically heteroscedastic: their sampling variance scales with the square of the true spectral power, which typically varies as a function of frequency. This probability model is the basis for the Gamma probability model used in our spectral decomposition framework.

## 3 Methods: Spectral decomposition as an estimation problem

### 3.1 Empirical ECoG data and ethics

We analyzed previously collected, publicly available electrocorticography (ECoG) recordings from the Neurotycho *Macaca fuscata* monkey dataset (M1). The original experimental recording protocol is described in (Yanagawa et al., 2013), and the surgical implantation procedures and electrode coverage are described in (Nagasaka et al., 2011). No new animal experiments were performed for the present study.

The original data collection and surgical procedures were approved by the RIKEN ethics committee under protocol No. H24-2-203(4) and were conducted in accordance with the recommendations of the Weatherall report, “The use of non-human primates in research.” The implantation surgery was performed under sodium pentobarbital anesthesia, efforts were made to minimize animal suffering, and no animal was sacrificed. Because the present study is a secondary analysis of previously collected non-human-primate data, informed consent was not applicable.

### 3.2 Overview

Our goal is to study how different spectral decomposition methods perform when applied to the same data. For example, Fig. 1A shows a 3-second window of ECoG recorded from a *macaque* undergoing propofol anesthesia. When `specparam` and our proposed method `SL_specdecomp` are used to decompose the power spectrum of this window, they estimate different rhythmic and broadband patterns (Fig. 1B–C). For example, `specparam` attributes the power below 2 Hz mostly to the broadband component, while `SL_specdecomp` attributes it mostly to a rhythm. These differences are also apparent in broadband biomarkers such as the spectral slope. Fig. 1D shows the distribution of estimated 40–60 Hz log–log slope across time windows: `specparam` systematically estimates flatter slopes, with more variability across windows than `SL_specdecomp`. Because the ground-truth spectra are not available *in vivo*, however, Fig. 1A–D alone cannot determine which model more accurately attributes spectral power to the underlying generative components.

**Figure 1.**
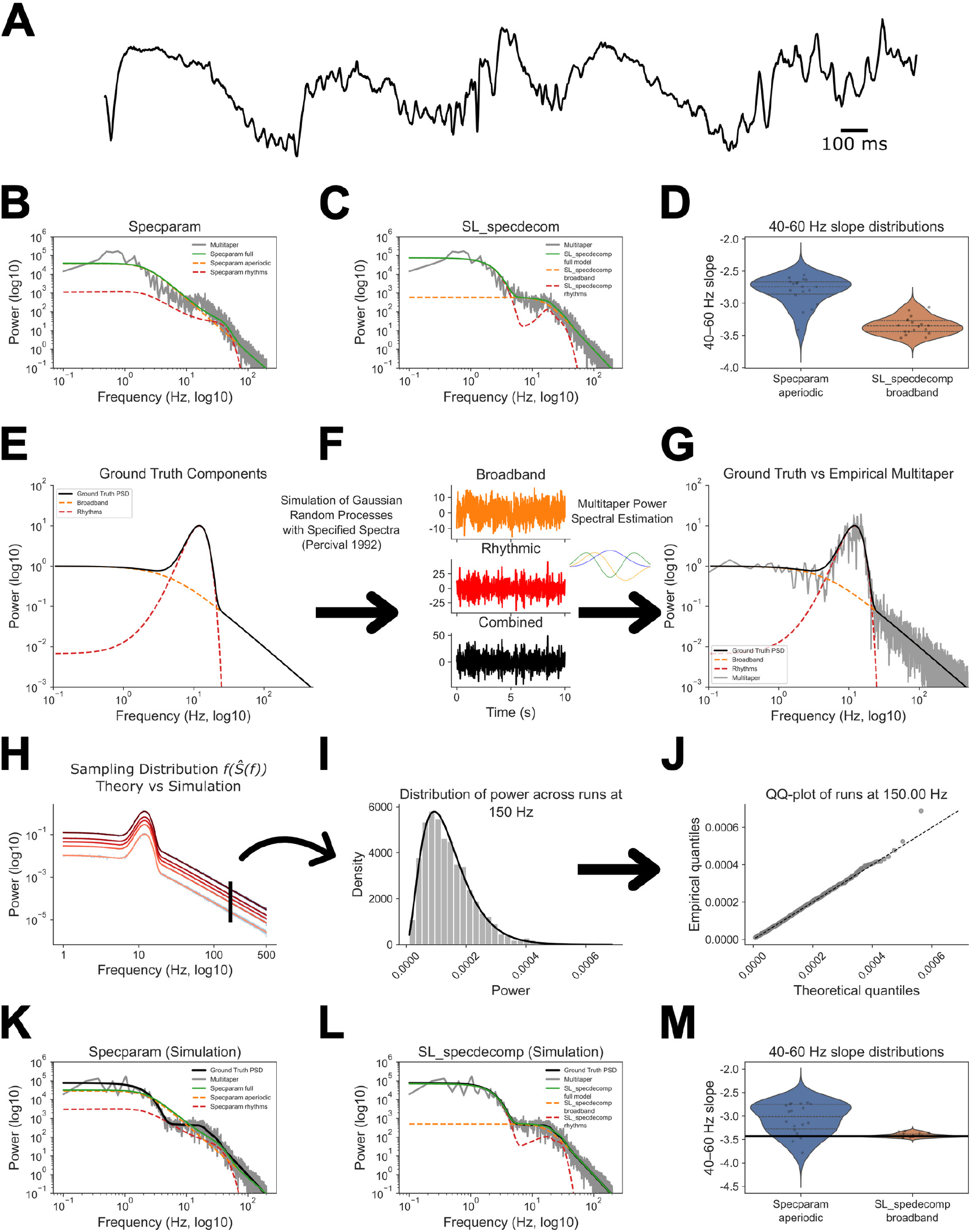
Different spectral decomposition methods yield meaningfully different decompositions of the same data, so we compare them using simulation. Real data: (A) Example macaque ECoG trace during propofol anesthesia showing slow-waves with a prominent alpha rhythm. (B–C) The multitaper spectral estimate of a representative window decomposed by specparam versus SL_specdecomp, illustrating different decompositions of rhythmic vs broadband power. (D) 40–60 Hz slope distributions across windows differ between the two spectral decomposition methods, but ground truth is unavailable *in vivo*. **Simulations:** (E–G) Realistic simulation pipeline: specify a ground-truth spectrum with broadband + rhythmic components (E), simulate Gaussian-process time series and estimate multitaper spectra (F), and compare ground truth to multitaper estimates (G). (H) shows the per-frequency empirical sampling distribution (percentiles) alongside the theoretical Gamma distribution; (I) illustrates the histogram at a single example frequency (150 Hz) compared to the theoretical Gamma PDF; and (J) shows the Q–Q plot at the same frequency, demonstrating that the empirical and theoretical quantiles align. (K–M) Under known ground truth, specparam misidentifies the slow rhythm as broadband power in this example (K) and shows biased/variable slope estimates across simulations (M). SL_specdecomp remains consistent with the true broadband trend (L) and concentrates near the true 40–60 Hz slope (M, black line) across simulations.

We therefore use simulations of Gaussian processes with known ground truth to evaluate when and how model assumptions drive estimation error. We study two common forms of model misspecification: (1) multiplicative (as opposed to additive) interactions between components in modeling the true power spectrum, and (2) a homoskedastic Gaussian (as opposed to Gamma) probability model. Our approach is: (i) define a theoretical power spectrum *S*_true_(*f* ) using either an additive or multiplicative functional form containing broadband and narrowband components; (ii) simulate a time-domain signal from *S*_true_(*f* ) and estimate its multitaper spectrum 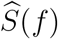; (iii) perform spectral decomposition by fitting models that use a correctly or incorrectly specified (a) functional form for the true spectrum and (b) probability model; and (iv) evaluate recovery of the ground-truth spectrum and derived spectral biomarkers. After isolating these misspecification effects in controlled settings, we perform a direct comparison of `specparam` and `SL_specdecomp` on both simulated and real data.

### 3.3 Generating simulated data based on ground-truth power spectra

#### 3.3.1 Defining ground-truth theoretical spectra

We define two-sided theoretical spectra *S*_true_(*f* ) that are symmetric in *f* (as required for signals that are real-valued in the time-domain) and explicitly exclude the zero frequency component (we set *S*_true_(0) = 0, corresponding to the assumption that the signal has mean zero). Rhythmic structure is represented using mirrored Gaussian bumps:

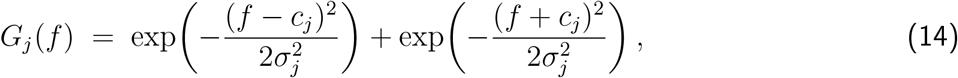

where *c*_*j*_ > 0 is the center frequency and *σ*_*j*_ > 0 is the bandwidth (standard deviation in Hz). For one-sided spectra we evaluate *G*_*j*_ on *f* > 0 but keep the mirrored form to account for the two-sided symmetry in the power spectrum of a real-valued signal (this is important for low-frequency rhythms, where the negative peak can spread to positive frequencies).

We model broadband structure using a generalized Lorentzian function:

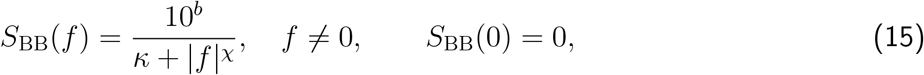

with broadband offset *b* (in log_10_ units), slope exponent *χ* > 0, and knee parameter *κ* > 0.

The *Additive Ground Truth* model adds the components:

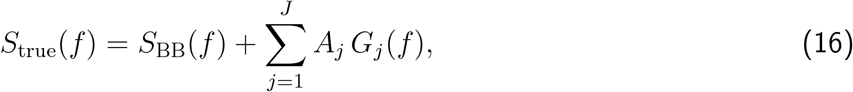

where *A*_*j*_ > 0 are linear-power amplitudes.

The *Multiplicative Ground Truth* model multiplies the components:

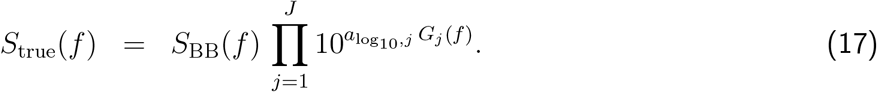

The same model can be described on a log scale:

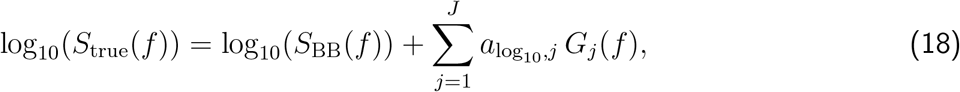

where 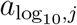 are peak amplitudes in log_10_-power units. Note that the multiplicative model (unlike the additive model) uses the Gaussian bumps in the exponent, so they will look more Gaussian-like when plotted on a log scale than a linear scale. We defined the model in this way to make the multiplicative model consistent with `specparam`.

The specific parameters that we used for our model comparisons below are described in Supplementary Section 3, especially Supplementary Tables 2 and 3.

#### 3.3.2 Simulating time-domain signals

Given *S*_true_(*f* ), we simulate a stationary mean-zero Gaussian process whose true spectrum equals *S*_true_(*f* ). Generating time series from a Gaussian process with a specified power spectrum is non-trivial. For example, it is not sufficient to simulate Fourier coefficients from Eq. (8) and apply an inverse Fourier transform. We use a method from (D. Percival, 1992), described in Supplementary Section 1.

#### 3.3.3 Estimating power spectra

We compute multitaper power spectra 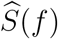 using *n*_tapers_ orthogonal Slepian tapers. Because multitaper estimates at nearby frequencies are correlated due to smoothing and finite spectral resolution, we subsample the positive-frequency axis (excluding *f* = 0) and evaluate 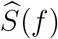 on a grid 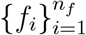 spaced by at least the multitaper spectral resolution. This yields frequency bins that are approximately independent.

### 3.4 Comparing constrained spectral decomposition models

To illustrate the consequences of a misspecified probability model alone, we evaluate the ability of several simple models to estimate the broadband height parameter (*b* or 10^*b*^, depending on the scale), when there are no rhythms. Under these circumstances, the additive and multiplicative ground truth models are equivalent.

Let *µ*_*i*_ ≡ *S*_model_(*f*_*i*_) denote the modeled true spectrum evaluated on the same grid. As a vector, denote it 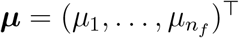. All spectral decompositions tested here assume that the broadband shape is known, so there is only one unknown coefficient *β* ∈ ℝ capturing the broadband height, either on a linear or a log scale. In other words:

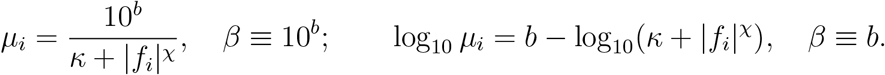

We study four model classes that isolate the effects of the probability model using a linear or a log scale:

1. **Gamma GLM with identity link (correct distribution; linear scale)**.

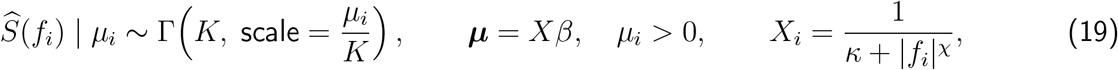

where *β* is interpreted as 10^*b*^.
2. **Gamma GLM with log link (correct distribution; log scale)**.

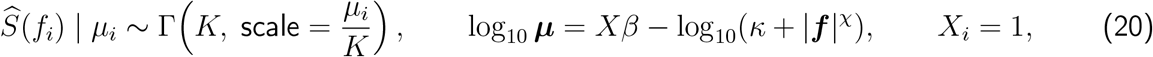

where *β* is interpreted as *b*.
3. **Gaussian OLS in linear power (misspecified distribution; linear scale)**.

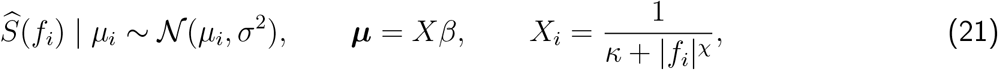

where *β* is interpreted as 10^*b*^.
4. **Log-transformed OLS (misspecified distribution; log scale)**.

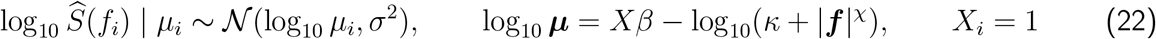

where *β* is interpreted as *b*.

Both of the Gamma GLM models are correctly specified with respect to the ground truth model, because (1) they have the correct probability model and (2) the additive vs multiplicative true spectrum models apply to how components combine, and these models only have one component. Both of the Gaussian models are misspecified with respect to the probability model (Gaussian instead of Gamma), and both assume that the variance is constant across frequencies. The difference is the scale at which the Gaussian noise is applied: linear vs log-transformed.

We evaluate models by their ability to recover (i) the ground-truth spectrum *S*_true_(*f*_*i*_) and (ii) the model-predicted high-*γ* mean power in 80–180 Hz.

### 3.5 Comparing SL_specdecomp **and** specparam

To evaluate the consequences of model misspecification under more general circumstances, we propose our estimation model, `SL_specdecomp`, and compare it to `specparam`.

#### 3.5.1 SL_specdecomp structure

`SL_specdecomp` uses a Bayesian framework, where the observed power spectrum is modeled with a Gamma likelihood and priors are placed on the rhythmic and broadband parameters. To model the true spectrum, we allow either additive or multiplicative structure. For the additive variant,

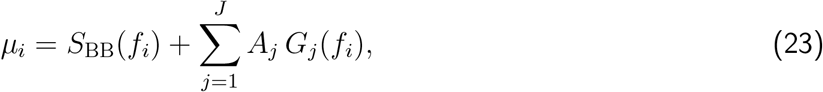

and for the multiplicative variant,

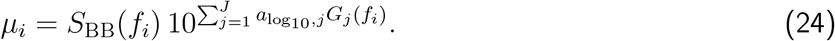

Both are paired with

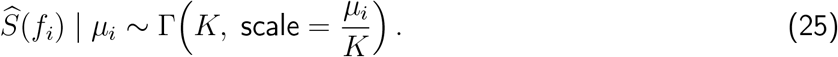

Note the correspondence of these equations to theory: Eq. (23) is based on Eq. (16); Eq. (24) is based on Eq. (17); and Eq. (25) is based on Eq. (12). The Gamma shape parameter is *K*, the number of uncorrelated direct estimates being averaged. In analyses where the spectrum is averaged only across independent tapers, *K* = *n*_tapers_; more generally, *K* can encode the effective number of uncorrelated direct estimates contributing to the averaged spectrum.

Let Θ denote the full parameter set. For the additive `SL_specdecomp` model,

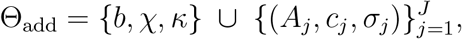

and for the multiplicative `SL_specdecomp` model,

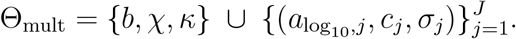

We place weakly informative priors on Θ and approximate the posterior 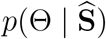 using NUTS (PyMC; optionally with a BlackJAX backend). Constraints are enforced by parameterization and prior support (e.g., *c*_*j*_ restricted to a specified frequency band, *σ*_*j*_ > 0 via truncated priors, and positive amplitudes via log-normal or half-normal priors; see Supplementary Section 2).

`SL_specdecomp` allows the user to choose the components and priors based on the experimental context. For example, in the propofol analyses below we used a slow-wave rhythm (0.1–4 Hz), an alpha rhythm (8–20 Hz), and a beta rhythm (20–30 Hz). We describe the settings that we used for Figures 3–8 in Supplementary Section 3, especially Supplementary Table 3.

#### 3.5.2 specparam structure

We use the `specparam 1.0` package (see Supplementary Section 3 for implementation details and fit settings) to perform spectral decomposition. `specparam` assumes a multiplicative structure in linear power (additive in log_10_ power) for the true spectrum and homoskedastic Gaussian residuals in log scale:

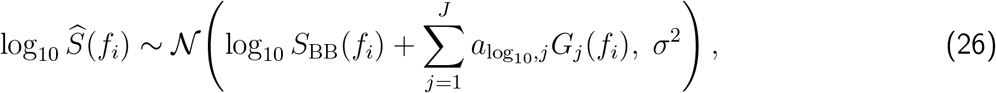

which implies the linear-power spectrum 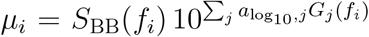. Note that when the data are generated according to the multiplicative true spectrum model in Eq. (17), `specparam` is correctly specified with respect to the structure of the true spectrum, and it is only misspecified with respect to the probability model.

Unlike `SL_specdecomp`, `specparam` uses an iterative algorithm to detect the number of rhythms and applies no constraints or priors on where they can occur. Here, we used the package to set the maximum number of rhythms equal to the ground truth number used for simulation, in order to partially account for this difference in flexibility between the two models.

#### 3.5.3 Model comparison

We compare `specparam` to `SL_specdecomp` on both simulated and empirical spectra. We evaluate models by their ability to recover (i) the ground-truth spectrum *S*_true_(*f*_*i*_) and (ii) derived spectral biomarkers, including recovered peak frequency, high-*γ* mean power in 80–180 Hz, and broadband slope on log_10_(power) vs log_10_(frequency) over 40–60 Hz. When cross-validated log-likelihood (CVLL) was evaluated, each 30 s window was partitioned into five non-overlapping 6 s folds. For each held-out fold, we averaged the multitaper spectra from the other four folds to obtain a training spectrum, fit each model to that training spectrum, and then evaluated the predictive log-likelihood of the held-out fold under the Gamma sampling model

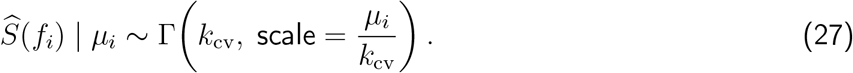

Thus, for model *m*, window *w*, and fold ℓ,

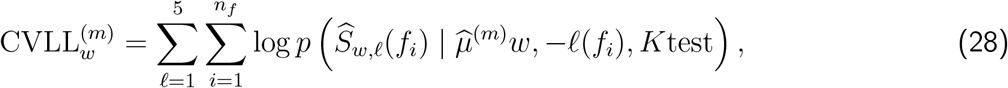

where 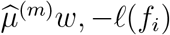 is the fitted mean spectrum from the training folds and *K*test = 1 for the held-out fold spectra. Higher CVLL indicates better out-of-sample predictive fit.

### 3.6 Package Implementation and Figure Code

All code for simulation, estimation, and figure generation is openly available. The spectral decomposition framework introduced here is implemented in the `SL_specdecomp` package (GitHub repository). Code used to generate the manuscript figures and analyses is available in Bloniasz_Stephen_Estimation. Gaussian-process simulations with analytically specified power spectra are implemented in `SL_GPsim` (GitHub repository).

## 4 Results

Widely used decomposition methods can estimate materially different broadband biomarkers from the same multitaper spectra, due to differences in their model structure and fitting approaches. Because ground truth is unavailable *in vivo*, we probe these discrepancies using a simulation regime that produces realistic spectra and multitaper sampling properties. We deliberately evaluate model performance in regimes where the ground-truth parameters are identifiable, so that errors can be attributed to model misspecification rather than to intrinsic non-identifiability of the decomposition problem. Here we will show that when the ground truth is known and identifiable, the most widely used spectral decomposition package (`specparam`) yields biased and unstable spectral decompositions, including confounding rhythms and broadband structure. Our proposed method (`SL_specdecomp`) yields more reliable decompositions in this regime.

### 4.1 Overview: Our approach to comparing spectral decomposition methods

We begin with an empirical motivating example from macaque ECoG during propofol anesthesia-induced unconsciousness (Fig. 1A), which exhibits prominent slow-wave structure alongside a narrowband “alpha” rhythm and “beta” rhythm. Applying two decomposition approaches to the same multitaper power spectrum—`specparam` versus `SL_specdecomp`—yields meaningfully different decompositions into rhythmic and broadband components (Fig. 1B,C): `SL_specdecomp` identifies the slow wave as a rhythm, while `specparam` attributes the low frequency power to the broadband component. These differences propagate to a standard broadband biomarker: the 40–60 Hz slope differs substantially between methods across analysis windows (Fig. 1D). Critically, because real data do not provide a ground-truth decomposition, it is not possible to determine from Fig. 1A–D alone which model is better able to capture the true underlying power spectral features.

To compare the accuracy of the decomposition methods, our approach is to construct a realistic simulation regime in which the true broadband and rhythmic components are known (Fig. 1E–G). We specify an analytic ground-truth spectrum with broadband and narrowband components (Fig. 1E), simulate Gaussian-process time series with this target spectrum (Fig. 1F), and estimate multitaper power spectra from the simulated time series. As expected, the observed power spectrum is a noisy realization of the ground-truth spectrum (Fig. 1G, gray and black lines, respectively). This provides a controlled setting that preserves key features of real spectra while enabling comparisons to ground truth.

A central statistical feature of power spectral estimates is their heteroscedastic sampling variability: the observed spectrum 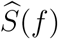 is Gamma distributed (Eq. (12)), so the variance of the power spectral estimate at a given frequency is proportional to the square of the true power spectrum. Across repeated simulations, the empirical sampling distribution of the estimated spectra closely matches this theoretical sampling distribution across frequencies (Fig. 1H). Fig. 1I focuses on a single frequency, showing that if you simulate the process many times, the distribution of power spectral estimates at 150 Hz (histogram) closely matches the theoretical Gamma distribution (black line). More formally, the Q-Q plot in Fig. 1J is focused on the 45-degree line, meaning that the theoretical and empirical distributions match. This motivates performing inference directly using Gamma likelihood, rather than assuming homoscedastic Gaussian residuals in linear or log-power space.

In a “simulated anesthesia” regime designed to have similar power spectra to the real data, `specparam` produces broadband distortions in example decompositions (Fig. 1K) relative to the true broadband trend. Here, the ground-truth spectrum has a slow wave, but `specparam` nevertheless attributes the low frequency power to the broadband component, as it did in the real data. In contrast, `SL_specdecomp` produces decompositions that remain consistent with the ground truth spectrum (Fig. 1L). Quantitatively, the 40–60 Hz slope—the same metric shown in real data in Fig. 1D—is biased under `specparam`, while `SL_specdecomp` remains concentrated around the true slope (Fig. 1M). Together, Fig. 1A–M motivates the remaining results by showing (i) that different spectral decomposition methods estimate different biomarkers in real recordings, and (ii) that realistic simulations with known ground truth can help suggest when such differences arise from misspecified modeling assumptions.

### 4.2 Result 1: A misspecified probability model leads to biased or variable broadband estimation in constrained decompositions

To disentangle the role of the probability model from the role of additive versus multiplicative true spectrum model, we studied models in which the spectral shape (broadband with no rhythm) was treated as known and only one coefficient (broadband height as a single scalar) was estimated. We compared two Gamma-GLMs (identity link vs. log link) to two Gaussian-based regressions (OLS in linear power and OLS in log-power; Fig. 2). In each case, the fitted model was evaluated on its ability to recover a broadband summary statistic (here, log_10_ power at 80 Hz) under repeated independent sampling (Fig. 2A,B).

**Figure 2.**
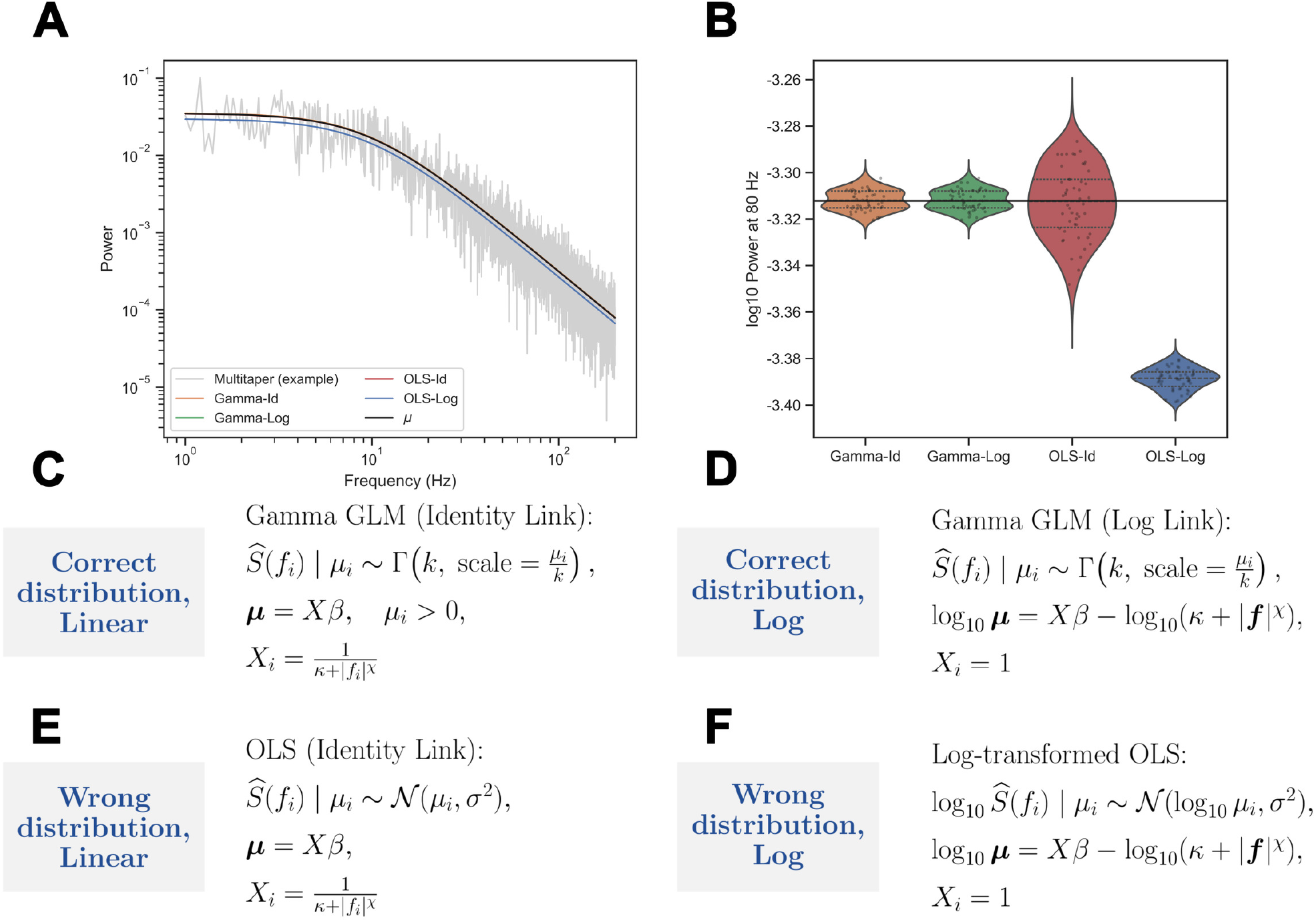
Model misspecification leads to biased/unstable estimates of broadband power, even when the spectral shape is known. With the broadband predictor shape treated as known, we estimate only a height coefficient and evaluate recovery of the power at 80 Hz (A–B, black line: ground truth). In this example, both Gamma-GLM with identity link (C) and Gamma-GLM with log-link (D) are correctly specified. Gaussian OLS (E) assumes the wrong sampling distribution for power spectra in linear power, while log-transformed OLS (F) applies a Gaussian least-squares objective after transforming the data to log-power. The two Gamma models are unbiased, while linear-power OLS is unstable and log-transformed OLS is biased when the fitted spectrum is interpreted on the original power scale (B).

Both Gamma-GLMs produced unbiased estimates with comparatively low variance (Fig. 2B), consistent with the correctly-specified probability model. Since there was only one component in the true spectrum, the different true spectrum models had no effect, so the two Gamma-GLMs were both correctly-specified. In contrast, OLS in linear power produced substantially higher variability, consistent with the fact that the linear-power residuals are strongly heteroscedastic under sampling (Fig. 2B, OLS-Id): in other words, the linear OLS model was unable to take advantage of the very low sampling variance at high frequencies. OLS in log-power produced a biased estimate of the power: while the log transform helps to stabilize the variance, the log-transformed residuals are asymmetric, with more extreme small values than large values, causing a negative bias in the height of the fitted spectrum. These comparisons demonstrate that misspecifying the probability model causes estimation methods to fail to recover the true broadband power, even when there is only one parameter to be estimated.

Thus, when the sampling distribution of power is modeled correctly, broadband height parameters can be recovered without systematic bias; in contrast, least-squares estimation under Gaussian residual assumptions yields bias and/or inflated variance, and these effects depend on whether fitting is performed on a linear or log scale for power. The consequences are limited in this case because the model fitting was extremely constrained: next we will evaluate how model misspecification can lead to unpredictable results when the model fitting is more flexible.

### 4.3 Result 2: In flexible decompositions, correctly specifying the probability model improves model performance, even when the true spectrum model is misspecified

We next considered the practical regime where decomposition methods estimate multiple parameters simultaneously (3 parameters for the broadband component (*b, χ, κ*) and 3 parameters for a rhythm). Using additive ground-truth simulations (Fig. 3A–C), we compared `specparam` to two Gamma-likelihood variants of `SL_specdecomp`: an additive true spectrum model (Eq. 23) and a multiplicative true spectrum model (Eq. 24). Because the different decompositions used different parameterizations of the true spectrum, we evaluated the models based on their ability to recover four summary features of the ground truth spectrum: rhythm height and peak frequency, high-*γ* mean power, and broadband slope (Fig. 3H-K). In representative fits, all methods tracked the overall spectrum on log–log axes (Fig. 3D–G), but differed in how power was decomposed near the rhythm and into higher-frequency broadband.

**Figure 3.**
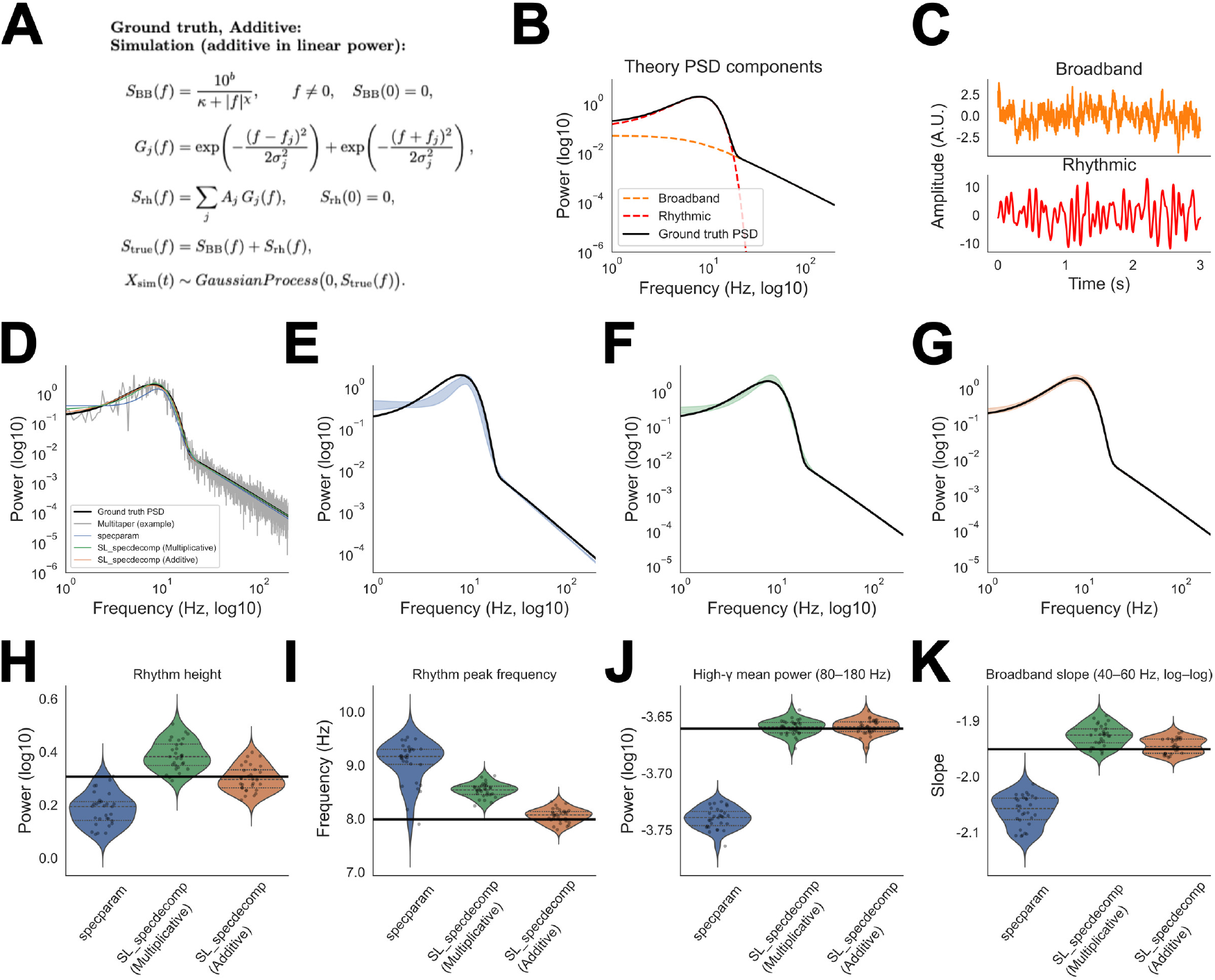
Additive ground truth: correctly specifying the true spectrum model and the probability model improves recovery of rhythmic and broadband biomarkers. (A) We simulate from an additive generative model, (B) where the ground truth spectrum has a broadband and rhythmic component. (C) An example simulated time series for each component. (D) Example fits to the multitaper spectrum of one simulated time series, comparing specparam to two Gamma-likelihood SL_specdecomp variants. (E) The 95 %ile distribution of fits is shaded for specparam. (F), (G) 95 %ile of fits for SL_specdecomp Gamma-multiplicative and SL_specdecomp Gamma-additive models, respectively. (H–K) Across repeated simulations, the Gamma-additive model is an unbiased estimator of rhythmic height, peak frequency (defined as the derivative-zero frequency), high-*γ* mean power (80–180 Hz), and broadband slope (40–60 Hz, log–log), with the ground-truth reference shown as the black line, while specparam exhibits systematic bias in peak and broadband features.

Across repeated simulations, `specparam` exhibited bias in all of the spectral features (Fig. 3H–K). The Gamma-multiplicative model (with misspecified true spectrum model but correctly specified probability model) was unbiased in estimating the high-*γ* mean power, and less biased than `specparam` for the other features. In contrast, the correctly-specified Gamma-additive model concentrated rhythm and broadband features near their ground-truth values (Fig. 3H–K). These results show that when the ground truth spectrum is additive, both correcting the probability model (Gamma) and using the correct spectrum model are necessary for stable recovery of peak and broadband biomarkers.

To isolate the role of the probability model in a regime favorable to `specparam`’s true spectrum model, we simulated spectra directly from a `specparam`-like multiplicative functional form (Fig. 4A–C). Even in this favorable setting, `specparam` still showed peak- and broadband-related estimation bias across repeated trials (Fig. 4H–K), indicating that getting the true spectrum model correct does not eliminate estimation error when the probability model is misspecified. Both Gamma-likelihood `SL_specdecomp` variants reduced these errors, with the correctly-specified Gamma-multiplicative model providing the most consistent recovery of rhythmic and broadband features near their ground-truth values (Fig. 4H–K). The remaining discrepancies between Gamma-additive and Gamma-multiplicative fits under multiplicative ground truth indicate that the true spectrum model still matters, but that correcting the probability model is a dominant driver of improved stability for power spectral estimates.

**Figure 4.**
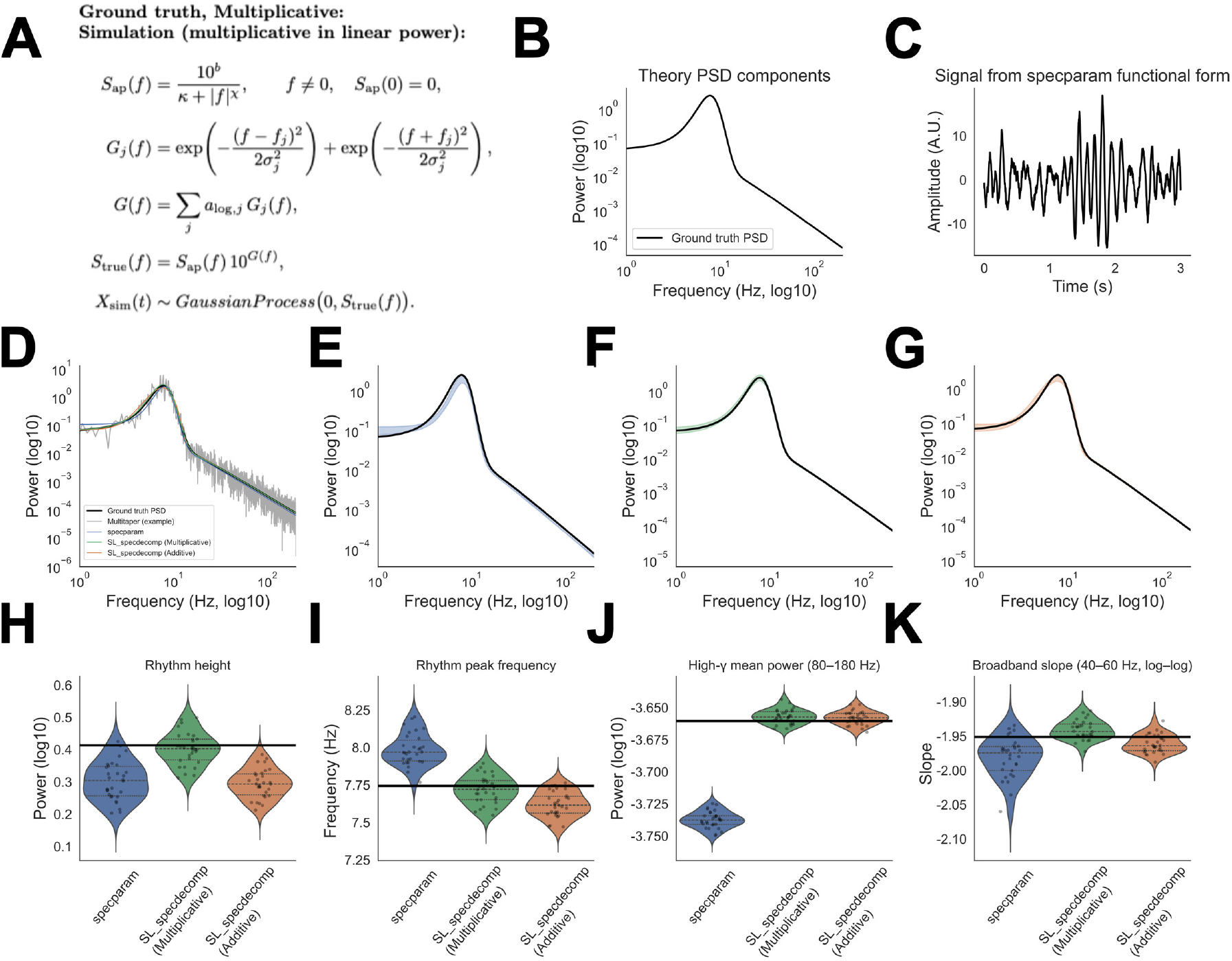
Multiplicative ground truth: correctly specifying the probability model remains influential for accurate decomposition. (A) We simulate from a multiplicative generative model (specparam-like), (B) the theoretical ground truth power spectrum, and (C) an example simulated time series. (D) Example fits to the multitaper spectrum of one simulated time series, comparing specparam to two Gamma-likelihood SL_specdecomp variants. The 95%ile distribution of fits is shaded for (E) specparam, (F) SL_specdecomp Gamma-multiplicative, and (G) SL_specdecomp Gamma-additive true spectrum model. (H–K) Under multiplicative ground truth, the Gamma-multiplicative model most consistently recovers rhythmic and broadband features, while specparam retains peak and broadband bias despite being matched to the generative mean structure.

### 4.4 Result 3: Increasing rhythm height induces rhythm–broadband confounding in specparam **but not in** SL_specdecomp

The previous simulations show that correcting the probability model improves recovery of broadband and rhythmic biomarkers. We next asked whether a strong rhythm can directly induce a spurious change in an inferred broadband biomarker even when the true broadband component is unchanged. To isolate this effect, we simulated spectra with a fixed additive broadband component and a single additive rhythm whose height was systematically increased across simulations (Fig. 5A). We then fit `specparam`, Gamma-multiplicative `SL_specdecomp`, and Gamma-additive `SL_specdecomp` to each simulated spectrum (Fig. 5B–D).

**Figure 5.**
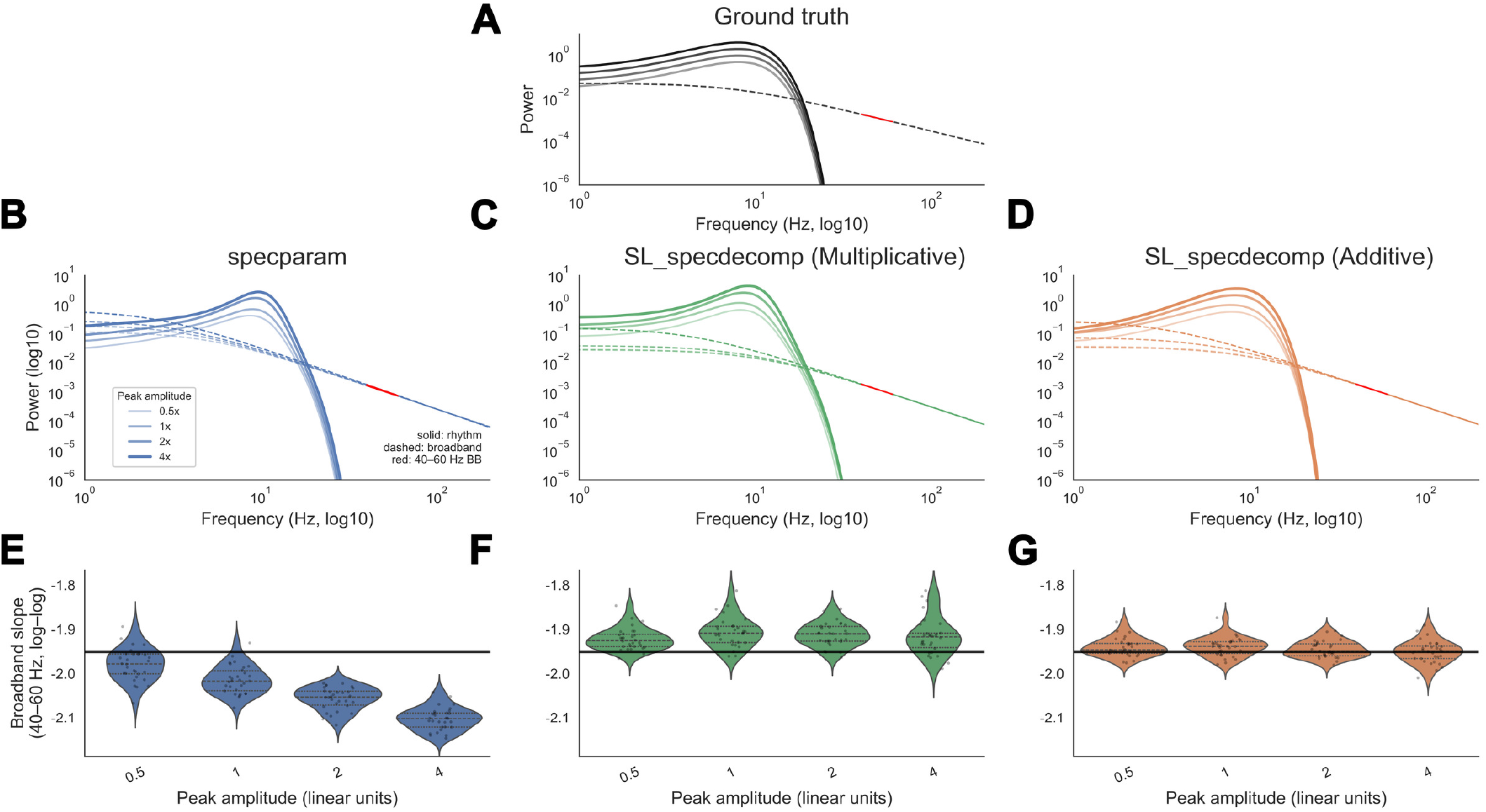
Increasing rhythm height induces rhythm–broadband confounding in specparam but not in SL_specdecomp. (A) Additive ground-truth spectra used for the rhythm-height sensitivity benchmark. The broad-band component is fixed across conditions, while the additive rhythmic component is varied across four amplitudes (0.5×, 1×, 2×, 4× ). The dashed curve shows the fixed broadband component, and the red segment marks the 40–60 Hz slope-evaluation band. (B–D) Representative fitted decompositions for specparam (B), Gamma-multiplicative SL_specdecomp (C), and Gamma-additive SL_specdecomp (D). Solid colored curves denote inferred rhythmic components, dashed colored curves denote inferred broadband components, and red segments mark the slope-evaluation band. (E–G) Sampling distributions of the inferred 40–60 Hz log–log broadband slope across repeated simulations as the ground-truth rhythm amplitude increases. The horizontal black line denotes the true slope of the fixed generating broadband component. specparam shows a rhythm-height-dependent shift in estimated broadband slope, even though the broadband component is unchanged. Gamma-multiplicative SL_specdecomp shows modest bias and variability but does not show the same systematic rhythm-height-dependent slope distortion. Gamma-additive SL_specdecomp remains concentrated near the true broadband slope across rhythm amplitudes.

As rhythmic height increased, all three decomposition methods struggled to capture the shape of the broadband component below about 10 Hz, i.e. the portion of the spectrum that is dominated by the rhythm (Fig. 5B–D). This reflects a known fundamental difficulty with spectral decomposition: in regions of the power spectrum where rhythms dominate, the data provide orders of magnitude less information about the broadband component (Bloniasz et al., 2025). In this simulation, the entire broadband component below the knee is dominated by a rhythm, providing very weak information to the models about the low frequency broadband shape. This is a situation that is likely to occur in real data, where low frequencies are often dominated by rhythms Buzsáki et al., 2012; Miller et al., 2009.

For `specparam`, this error in estimating the broadband component for low frequencies also produced a systematic shift in the inferred 40–60 Hz broadband slope, despite the fact that the ground-truth broadband slope was held fixed (Fig. 5E). This is rhythm–broadband confounding: a change in the fitted broadband biomarker is induced by a change in rhythmic amplitude rather than by any change in the generating broadband component. The effect is especially concerning because broadband slope is often interpreted physiologically; under this misspecified decomposition, a purely rhythmic manipulation can appear as a broadband change.

Gamma-additive `SL_specdecomp` did not show this confounding effect. Its slope estimates remained concentrated near the true broadband slope across rhythm heights (Fig. 5G), indicating that the broadband slope remains identifiable in this benchmark when the probability model and additive mean structure are aligned with the data-generating process. This is a direct result of the corrected probability model: under the Gamma model, the data has much more precision (lower variance) when the power is low, i.e. at higher frequencies. As a result, `SL_specdecomp` appropriately prioritizes the fit at high frequencies, and does not let the uncertainty at lower frequencies affect the high-frequency fit.

For the same reason, the Gamma-multiplicative `SL_specdecomp` variant also didn’t have confounding between the rhythmic height and broadband slope (Fig. 5F). It did have modest bias and variability, consistent with fitting a multiplicative peak structure to data generated by additive superposition, as seen in Result 2. However, this bias was not the same failure mode observed for `specparam`: increasing rhythm height did not induce the same systematic rhythm-height-dependent shift in the inferred broadband slope. Thus, the most damaging effect isolated here is not merely generic model bias, but rhythm-dependent distortion of a broadband biomarker under the `specparam` fitting criterion.

### 4.5 Result 4: We can compare model fits using cross-validated log-likelihood

One of the advantages of knowing that power spectra are Gamma distributed is that we can directly compare proposed (fitted) power spectra using their likelihood; i.e. better models will assign higher probability to the observed power spectrum. To account for possible overfitting, we use cross-validated log likelihood, which measures how well a fitted decomposition predicts held-out spectral observations. For each simulation trial, we computed CVLL for `specparam`, Gamma-`SL_specdecomp` with multiplicative true spectrum model, and Gamma-`SL_specdecomp` with additive true spectrum model (Fig. 6), and we ask whether the CVLL is consistently higher for the correctly-specified model versus the misspecified models.

**Figure 6.**
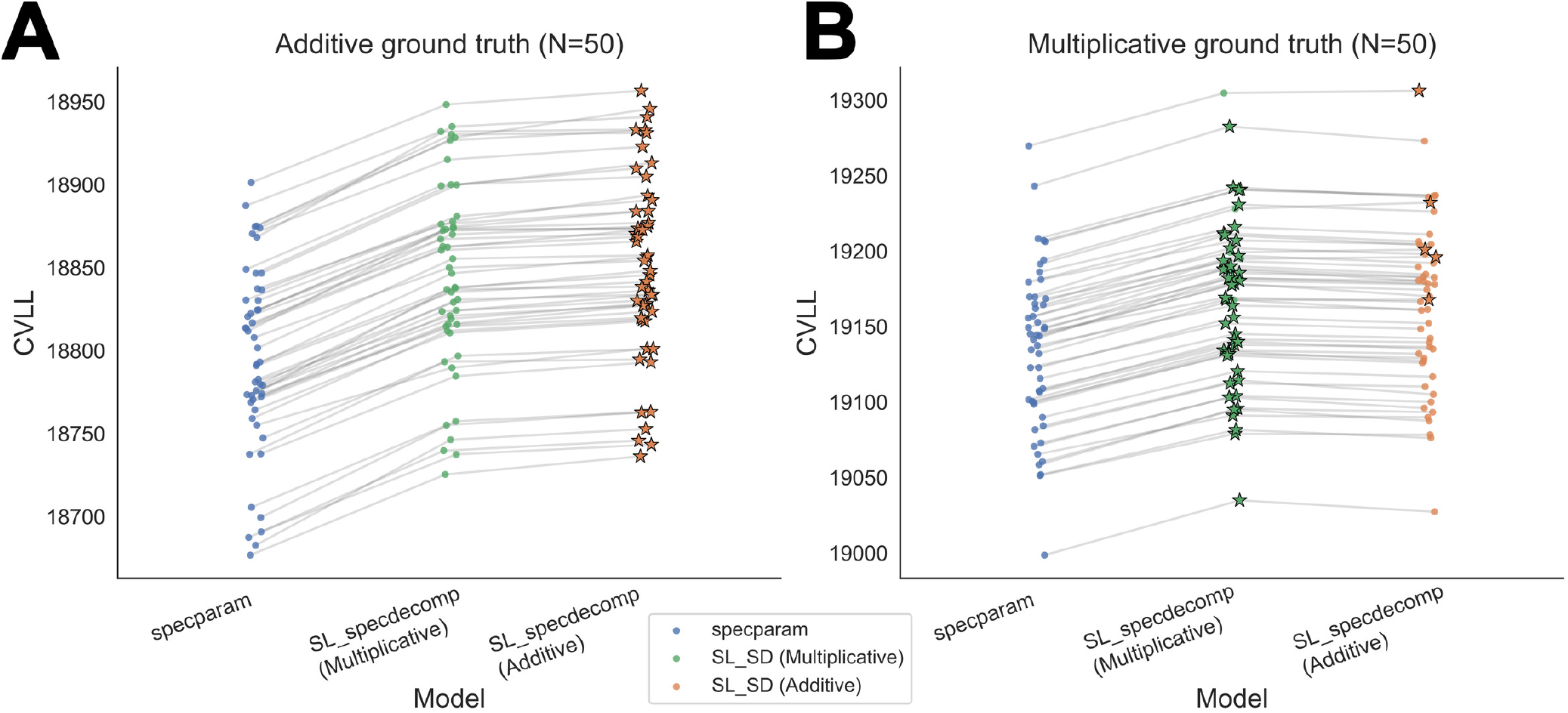
Cross-validated log-likelihood (CVLL) favors the correct true spectrum model and suggests Gamma-based models describe data better almost every time. For each simulated trial, we compute a trial-level CVLL score for each fitted decomposition and connect the three model scores with a line; star markers indicate the highest CVLL for each trial. (A) Under additive ground truth, the Gamma-additive model attains the highest CVLL in all trials. (B) Under multiplicative ground truth, the Gamma-multiplicative model attains the highest CVLL in most trials, while Gamma-additive models win for several trials.

Under additive ground truth (Fig. 6A), CVLL always ranked the Gamma-additive model highest, with the Gamma-multiplicative model intermediate and `specparam` lowest. Thus, when the data are generated by additive superposition in linear power, CVLL selects both the correct probability class (Gamma) and the correct true spectrum model (additive).

Under multiplicative ground truth (Fig. 6B), CVLL most often ranked the Gamma-multiplicative model highest, while the Gamma-additive model has several simulations marked as the highest CVLL, and `specparam` was always the lowest. Together, these results show that (i) a Gamma probability model yields systematically better predictive performance for power spectral estimates, and (ii) among Gamma-based models, CVLL discriminates additive versus multiplicative spectra in the direction of the true generative regime most of the time, suggesting that having a prior model for the generative mechanism is useful.

We do urge caution, however, in over-interpreting the comparison of the Gamma-additive and Gamma-multiplicative models here. Because the true spectrum models are different, the spaces of possible spectra that they can capture are different. For example, the rhythms in the multiplicative model will have Gaussian-like shapes when plotted on a log scale, while the rhythms in the additive model will have Gaussian-like shapes when plotted on a linear scale (and as a result, the tails will tend to be dominated by other spectral components). While we have shown here that CVLL can identify the differences between these cases in simulation, the result is driven by these differences in the fitted shape of the spectra, not whether the components actually compose additively versus multiplicatively. In real data, both of these are controversial questions: (1) what are the best shapes for power spectral components and (2) do they compose additively or multiplicatively (or both). We leave further study of these questions for future work, and move on to evaluate the comparison between `specparam` and just the Gamma-additive `SL_specdecomp` model below.

### 4.6 Result 5: In real macaque ECoG, specparam and SL_specdecomp produce systematically different rhythmic–broadband decompositions in both awake and anesthetized states

We next compared `specparam` to `SL_specdecomp` on real macaque ECoG during an awake resting period and during propofol anesthesia. Although both methods are fit to the same multitaper spectra, they can produce visibly different decompositions of power into broadband versus rhythmic structure. Figure 7 shows representative awake (panels A and B) and anesthetized (panels C and D) spectra estimated from 30 s windows, as decomposed by `specparam` and `SL_specdecomp`. In both the awake and anesthetized examples, the full fitted spectra can appear broadly similar on log–log axes, yet the decomposed broadband and rhythmic components differ substantially between methods, consistent with the misspecification effects isolated in simulation above.

**Figure 7.**
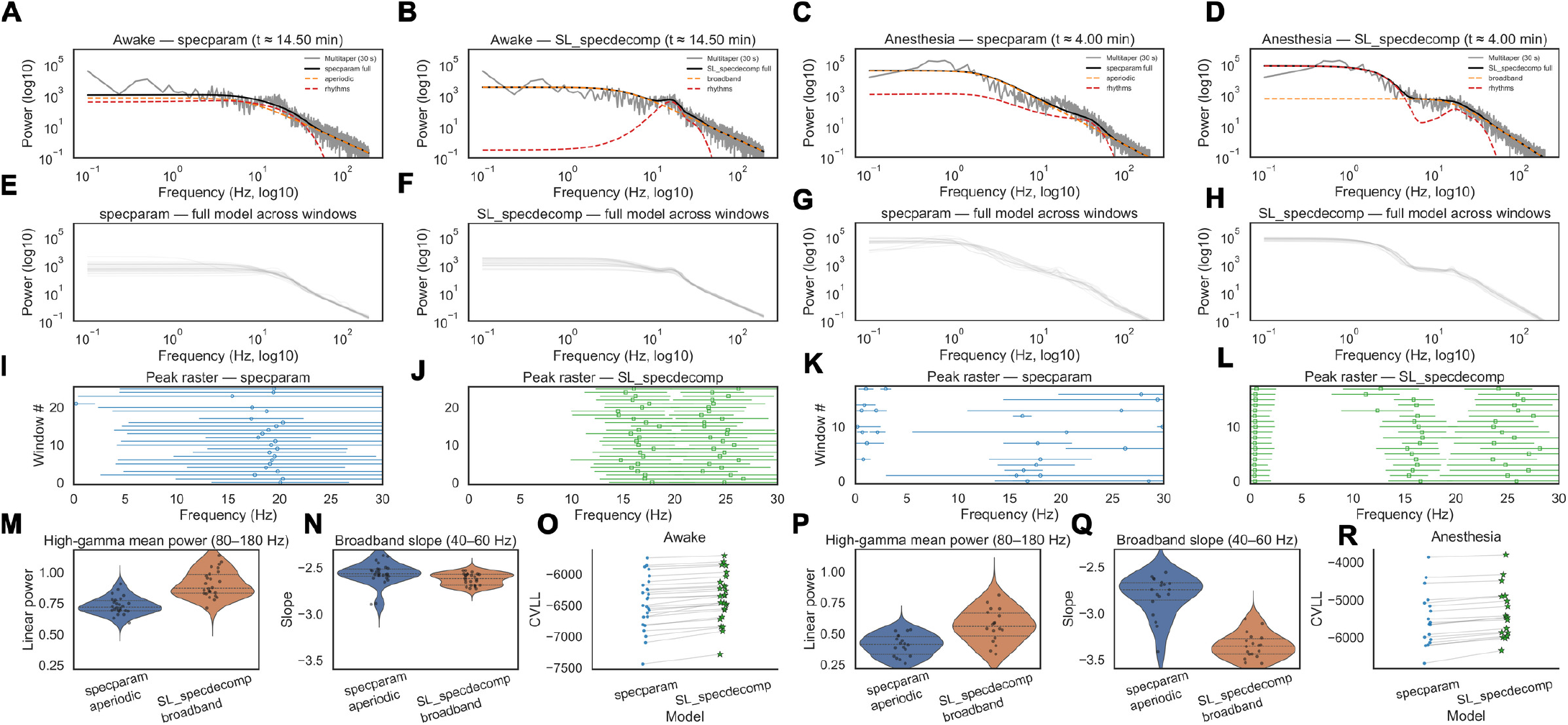
Real ECoG recordings: SL_specdecomp and specparam yield different decompositions in awake and propofol-anesthetized states. (A–D) Representative multitaper spectra from 30-s windows (gray) from macaque ECoG during an awake resting window (A,B) and an anesthetized window (C,D), decomposed by specparam (A,C) and by SL_specdecomp (B,D; Gamma-additive). Black curves denote each method’s full fitted spectrum; dashed curves show the inferred broadband component and rhythmic contribution. (E–H) Full fitted spectra across all analyzed windows within each state, highlighting systematic differences in the implied broadband trend across windows between methods. (I–L) Peak rasters summarizing the fitted rhythmic components across windows: markers denote peak center frequency and horizontal bars denote the corresponding bandwidth (center ± FWHM/2). Compared to SL_specdecomp, specparam produces a qualitatively different peak map in both states. (M,N,P,Q) Across windows, features of the inferred broadband component differ systematically between methods, including high-*γ* mean power (80–180 Hz; linear power) and log–log broadband slope (40–60 Hz). In particular, SL_specdecomp shows a markedly steeper broadband slope in anesthesia than in wakefulness, whereas specparam shows a smaller state shift. (O,R) Cross-validated log-likelihood (CVLL) favors SL_specdecomp across windows in both awake and anesthetized states, indicating improved predictive accuracy for the observed spectra. The star markers indicate the higher-CVLL model for each window.

These differences are systematic across time windows within each state. When we overlay the fitted full spectra across all windows (Fig. 7E–H), `specparam` and `SL_specdecomp` yield different implied broadband shapes across frequencies, which extends to derived broadband features. Across windows, `SL_specdecomp` yields systematically shifted estimates of high-*γ* mean power (80–180 Hz) and 40–60 Hz log–log slope relative to `specparam` (Fig. 7M,N,P,Q). Because it is extremely common to interpret these features physiologically (e.g., as measures of population firing-rates or excitation/inhibition balance), the fact that different methods yield different decompositions could lead to significant differences in interpretation. For example, `SL_specdecomp` predicts that the broadband slope becomes substantially steeper (more negative) during anesthesia relative to the awake state, whereas `specparam` shows a much smaller state difference. For context, Supplementary Fig. 1 shows the estimated slopes across time during the full recording session and marks the windows used for the representative Fig. 7 panel fits.

With respect to rhythms (Fig. 7I–L), `specparam` has more variability across time windows than `SL_specdecomp`. This is a direct result of the fact that `specparam` is more flexible: it allows rhythms to occur at any frequency, and adds rhythms as needed up to a maximum (here, 2 for awake and 3 for anesthesia). `SL_specdecomp` in its current form is constrained to have a prespecified set of rhythms, each in a known band (here, 8–20 Hz and 20–30 Hz for awake and 0.1–4 Hz, 8–20 Hz, and 20–30 Hz for anesthesia). We expect that this constraint will be helpful in some situations but not in others, so future versions of `SL_specdecomp` will have options for more flexible rhythms.

Finally, cross-validated log likelihood favors `SL_specdecomp` in both awake and anesthetized states (Fig. 7O,R), indicating that the Gamma-based model fit is more consistent with held-out spectral observations. Because the 20–30 Hz beta rhythm is not always included in models of propofol-induced unconsciousness, we also show in Supplementary Fig. 2 that CVLL favors including the beta rhythm during 66.7% of anesthetized windows for the `SL_specdecomp` model. During the awake state, in contrast, CVLL favors the model without the 20–30 Hz beta rhythm in 85.7% of windows. Together, these real-data results mirror our simulation findings: different modeling choices can induce systematic differences in broadband and rhythmic biomarkers even when applied to identical observed spectra.

### 4.7 Result 6: In real-window simulations, SL_specdecomp is closer to ground truth than specparam

Result 5 established that on real ECoG data, `specparam` and `SL_specdecomp` produce materially different broadband biomarkers from the same multitaper spectra, but real data do not provide ground truth. Here we convert that real-data comparison into a controlled test by generating synthetic datasets that match the spectrum of the real example time window with known ground-truth parameters.

For each state (awake and propofol anesthesia), we select a representative 30 s window from the real analysis (as in Result 5) and simulate a dataset with the same number of 30 s windows as the real study. We then fit `specparam` and `SL_specdecomp` to each synthetic window and evaluate model performance using cross-validated log-likelihood (CVLL). Fig. 8 has the same layout as Fig. 7, but for this simulated data.

**Figure 8.**
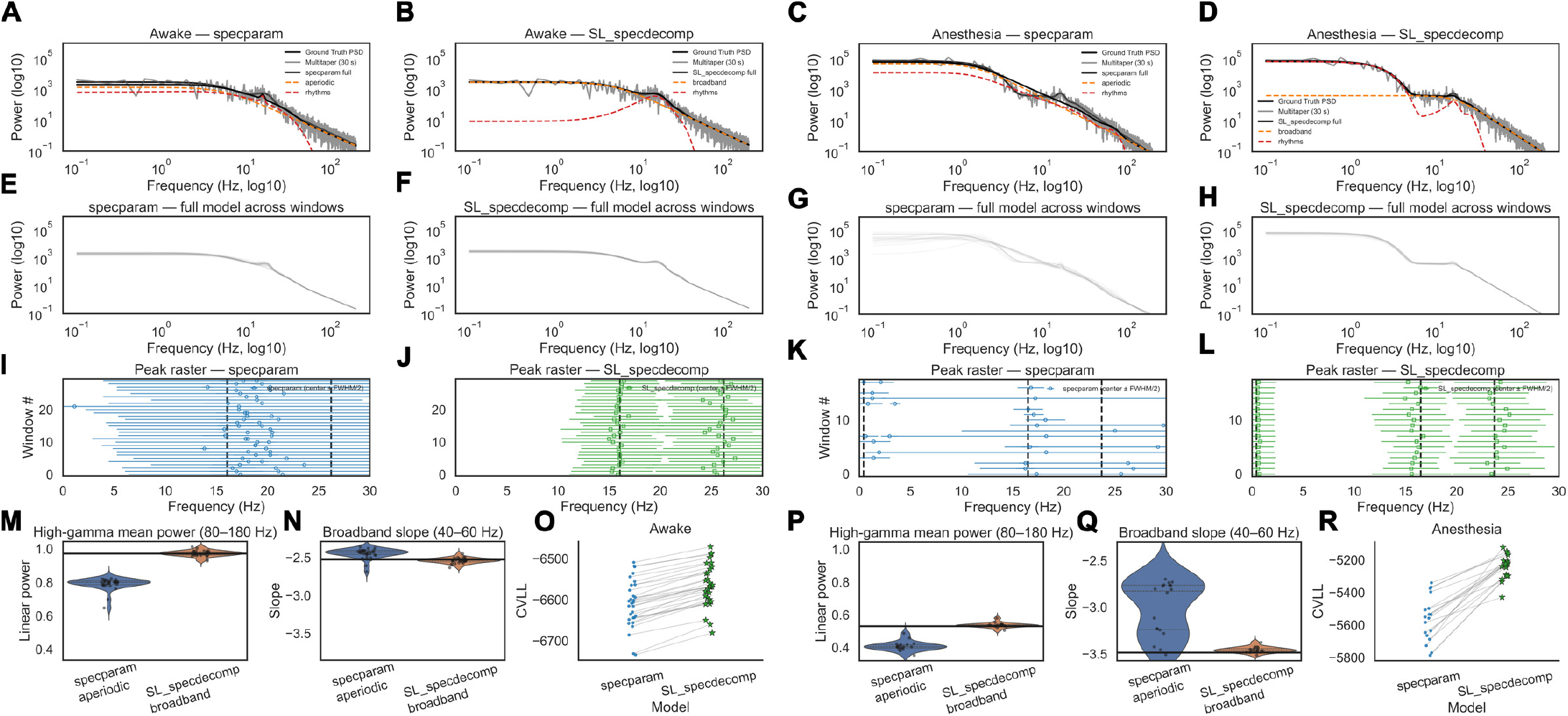
Ground-truth simulation generated from a representative real ECoG window shows that SL_specdecomp is closer to ground truth and that specparam errors are often rhythm-driven. For each state (awake and propofol anesthesia), we select one representative 30 s real-data window (as in Result 5) and use the corresponding fitted spectrum parameters to generate independent 30 s synthetic windows with the same state-specific full durations used in the analysis. (A–D) Example decompositions for awake (A,B) and anesthesia (C,D). Gray curves show multitaper power; thick black curves show the ground-truth spectrum used to generate the data. Colored curves denote each method’s fitted full spectrum (solid) and its inferred broadband and rhythmic components (dashed). Despite similar full-spectrum fits, specparam and SL_specdecomp produce different decompositions into broadband versus rhythmic components. (E–H) Fitted full spectra across all simulated windows, illustrating that method differences persist even when independent windows are generated from identical ground-truth parameters (i.e., absent real-window heterogeneity). (I–L) Peak rasters across windows (center frequency with horizontal bars indicating bandwidth, center ± FWHM/2), showing that specparam’s inferred peak structure differs systematically from the generative structure. (M,N,P,Q) Broadband features computed from the inferred broadband component: high-*γ* mean power (80–180 Hz; linear power) and log–log slope (40–60 Hz). Black horizontal lines denote the ground-truth values. In panels M and P, specparam underestimates high-*γ* mean power relative to both ground truth and SL_specdecomp. In panels N and Q, specparam estimates higher (less negative) broadband slopes, whereas SL_specdecomp remains closer to the true values. (O,R) Cross-validated log-likelihood (CVLL) favors SL_specdecomp across simulated windows in both states, indicating improved predictive accuracy under the correct sampling model. The star markers indicate the higher-CVLL model for each simulated window.

The performance of the models on simulated data is remarkably similar to their performance on the real data, suggesting that the simulation is well-constructed to mimic real data and study the sources of the differences in model fits.

In the representative-window panels (Fig. 8A–D), both methods can approximately track the observed multitaper spectrum, yet they produce different decompositions of power into broadband versus rhythmic components. Because the ground-truth spectrum is known in this simulation, we can attribute these discrepancies directly to model error: `specparam`’s Gaussian-in-log-power residual model often fails to identify the true rhythms and misattributes power to rhythmic versus broadband sources, whereas `SL_specdecomp`’s Gamma likelihood on linear power with additive true spectrum model more faithfully recovers the ground-truth rhythmic and broadband components.

Similar to the real data results, `specparam`’s rhythms are more variable (Fig. 8I-L), its measures of high-*γ* power are lower (Fig. 8M, P), and its measures of broadband slope are higher (Fig. 8N, Q) than `SL_specdecomp`. Here, however, we can compare the fits to the ground truth, showing that `SL_specdecomp` is systematically closer to the true values. In the anesthesia simulation in particular, `specparam` frequently fails to detect the true slow rhythm and instead attributes power below 1 Hz to the broadband component (Fig. 8K), which causes it to misestimate the overall shape of the power spectrum (Fig. 8G) similar to the real data (Fig. 7C, G, K).

These simulations recapitulate the failures described in Results 1-3: `specparam` systematically under-estimates broadband height (Result 1), fails to recover true rhythmic and broadband spectral features (Result 2), and shifts power between rhythmic and broadband components (Result 3). In addition, we see here that the flexibility of `specparam` can cause brittleness, in that it gives very different qualitative fits (number of rhythms, etc) to data generated from the same underlying model.

Finally, CVLL favors `SL_specdecomp` across simulated windows in both states (Fig. 8O,R), confirming the result seen in the real ECoG example. Together, these simulations indicate that the differences between `specparam` and `SL_specdecomp` observed in real ECoG are well explained by `specparam`’s misspecified modeling assumptions.

## 5 Discussion

Power spectral decomposition is now a routine step in analyzing neural field potential recordings, and the resulting parameters are frequently interpreted as biomarkers of network physiology and brain state (Donoghue, 2025; Donoghue & Watrous, 2023; Donoghue et al., 2020). Here we show that two modeling assumptions underlying many popular decomposition approaches are systematically misspecified: (i) the true spectrum is often assumed to be multiplicative in linear power (additive in log-power), despite the additive superposition implied by standard biophysical models of field potentials; and (ii) the residual/probability model is often treated as Gaussian and homoscedastic, despite the Gamma sampling distribution and intrinsic heteroscedasticity of direct spectral estimators. Together, these misspecifications can induce systematic estimation error and unstable decomposition of power into rhythmic versus broadband components, which in turn can lead to misinterpretation of widely used broadband biomarkers.

### 5.1 Insights and takeaways

Our analyses emphasize that power spectral decomposition is fundamentally an estimation problem, and attention to how model misspecifications affect estimation and interpretation is essential. First, several widely used methods (including `specparam`) implement a model that is additive in log-power, thereby imposing a multiplicative relationship between broadband background and rhythmic peaks (Donoghue et al., 2020). This assumption is convenient computationally but is not naturally aligned with the standard linear superposition model of field potentials, in which the recorded voltage is a sum of many current generators and the corresponding spectral power is additively decomposable into components (Eq. 1–3) (Bédard et al., 2004; Pettersen & Einevoll, 2008). An additive relationship between broadband and narrowband components is also consistent with the filtered point process model of local field potentials (Bédard et al., 2006; Bloniasz et al., 2025; Destexhe & Bédard, 2022). Second, power spectra derived from Gaussian time-domain processes via direct estimators are Gamma distributed with variance proportional to squared power (Eq. 12–13) (Kass et al., 2014; Kramer & Eden, 2016; D. B. Percival & Walden, 1993). Nonetheless, many decomposition pipelines optimize least-squares objective functions in linear power or log-power, which implicitly assume Gaussian residuals with frequency-independent variance (Donoghue et al., 2020; Medrano et al., 2025). These assumptions are incompatible with the actual sampling distribution of the estimators and therefore induce bias in the decompositions.

A central result is that misspecification manifests as systematic parameter misestimation even under well-controlled conditions. In fixed-shape GLM experiments (Result 1), correcting only the probability model (Gamma vs Gaussian) was sufficient to eliminate bias and stabilize broadband estimation. In full decomposition and rhythm-height simulations (Results 2–4), an incorrect probability model and/or true spectrum model (with `specparam`) produced distortions in both rhythmic parameters (e.g., peak height and peak frequency) and broadband summaries (e.g., high-*γ* mean power and log–log slope), while correcting one or both of these issues (with `SL_specdecomp` additive and multiplicative models) led to reduced bias and variance. Importantly, model misspecification translated into meaningfully biased biomarker distributions relative to known ground truth, as detected by the CVLL metrics. This distinction matters because many downstream physiological inferences are based on the estimated components.

One striking implication of our findings is that `specparam` can exhibit bias even when the true functional form is favorable to it. Under multiplicative ground truth (Result 2, Fig. 4), `specparam` is, in principle, matched to the functional form of the data-generating spectrum; nonetheless, it retains systematic peak-associated distortions and broadband bias. This indicates that the dominant source of error is not the parametric forms of the broadband component or the rhythmic peaks, but the mismatch between the Gaussian, homoscedastic residual model in log-power and the Gamma, heteroscedastic sampling distribution of direct spectral estimates. In this sense, the bias is a property of the estimation criterion: least-squares optimization in log space reweights errors across frequencies in a manner that does not correspond to the sampling variability of the estimator. This is specifically apparent at high frequencies, which is of great concern in estimation pipelines, because spectral decomposition is often used to accurately decompose these higher frequencies. This finding applies more broadly than `specparam`: any approach that fits parametric models to direct spectral estimates using Gaussian, homoscedastic residual assumptions (either in linear power or in a transformed space) inherits related distortions.

The rhythm-height sensitivity analysis makes the impact of the misspecification especially clear (Result 3). When the broadband component was fixed and only rhythmic amplitude was increased, `specparam` estimated a changing broadband slope. Thus, a purely rhythmic manipulation generated an apparent broadband effect. This is a critical interpretive risk for spectral decomposition in neuroscience: if a method reallocates power between rhythmic peaks and the broadband component as rhythm strength changes, researchers can mistake a change in network resonance (rhythms) for a change in asynchronous population activity or intrinsic timescales (broadband). `SL_specdecomp` avoided this confounding in the same benchmark, for both the additive and multiplicative variants. This result matters because generic bias and rhythm-broadband confounding are not equivalent: the former may only lead to errors in the magnitude of effects, while the latter can lead to fundamental misunderstandings of the underlying system.

Our macaque ECoG application offers another example of how poor model fits can change the interpretation of brain states. Recent naturalistic human ECoG/iEEG studies have used spectral decomposition to track brain state changes in real time (Alasfour & Gilja, 2024; Ervin et al., 2025; Zhu et al., 2024), arguing that brain state switches between distinct physiological modes. Our analysis of anesthetized data suggests that apparent switching can occur due to brittle model fits rather than true underlying changes in the power spectra. In particular, `specparam` frequently fails to detect the slow wave, instead capturing the low frequencies using the broadband component. This affects the slope of the broadband component for those windows, leading to highly variable spectra and broadband slopes (Fig. 7G,Q). This effect is even more apparent in the simulated data (Fig. 8G,Q), where `specparam` has a clear bimodal distribution in broadband slope due to switching between detecting the slow wave versus using the broadband to capture the slow wave. This apparent switching happens even though the ground-truth spectrum is the same for all windows. Under current practice, this pattern would be misinterpreted as switching between brain states with different excitatory-inhibitory balance. `SL_specdecomp`, in contrast, yields stable estimates of both the rhythmic and the broadband components (Fig. 7H,Q and Fig. 8H,Q). In both real and simulated data, `SL_specdecomp` would characterize anesthesia as having a consistent slow wave and relatively stable excitatory-inhibitory balance (with more inhibition than the awake state). Not only is this interpretation more consistent with scientific consensus on brain dynamics under propofol anesthesia – it also fits the data better, as we show with our proposed model comparison metric, CVLL (Fig. 7R and Fig. 8R).

This approach is another major contribution of our work: we show how CVLL can be used as an explicit model comparison tool, which can be used to test both alternative spectral decomposition approaches (Figs. 6–8) and physiological hypotheses (Supplementary Fig. 1). In simulation, we show that Gamma-likelihood models consistently attain higher CVLL (Results 2 and 4), and that the Gamma-additive model performs best across essentially all trials when the ground truth spectrum is additive (the biophysically motivated regime).

In real data, CVLL can be used to select between qualitatively different interpretations of the same brain state. For example, many studies report changes in broadband slope under anesthesia or altered consciousness and interpret these changes as alterations in excitation/inhibition balance or circuit time constants (Donoghue, 2025; Gao et al., 2017, 2020; Kramer & Chu, 2024). An alternative hypothesis is that apparent slope changes partly reflect a beta-band rhythm that emerges during sedation, sometimes described as “paradoxical excitation,” and thought to arise from beta-frequency network resonances (Ching et al., 2010; McCarthy et al., 2008). Our CVLL-based model-selection procedure provides a direct way to evaluate this possibility window by window, by comparing models with and without a beta rhythm. In the anesthetized state, the model comparison supports a beta-band rhythm in 66.7% of windows (Supplementary Fig. 1B; Figure 7K,L). Furthermore, both `specparam` and `SL_specdecomp` show a steepening of the spectral slope during anesthesia relative to the awake state (Figure 7 and Supplementary Fig. 1). In fact, `SL_specdecomp` estimates more spectral steepening than `specparam`, despite explicitly including a beta-band component in the anesthetized model. Physiologically, this could suggest that propofol causes both a change in the circuit time constants and a network resonance at beta frequencies.

Given that this is a single result in one monkey, it does not imply that every reported anesthesia-related slope change is a combination of rhythmic and broadband effects. Rather, it highlights that misspecified decompositions can attenuate or distort true broadband state differences when rhythms are present. When rhythmic changes co-occur with broadband shifts, decomposition methods must correctly attribute power to the appropriate components. Our simulations and real-data comparisons suggest that Gamma-likelihood inference in linear power is substantially more stable in this regime, making it a safer foundation for interpreting broadband biomarkers in the presence of strong rhythms.

Our results motivate caution when interpreting prior findings that depend critically on separating narrow-band and broadband components, particularly when the findings report changes in both rhythmic and broadband features. Because we view spectral decomposition as an estimation problem, design choices in the model specification will determine identifiability, interpretation, bias, and uncertainty quantification. We therefore recommend that future decomposition methods explicitly specify (i) the assumed sampling distribution of the spectral estimator, (ii) whether the true spectrum model is additive or multiplicative in linear power, and (iii) the scoring rule used for model comparison, ideally evaluated under a likelihood aligned with the estimator’s known distribution.

### 5.2 Limitations and future directions

Our study also highlights limitations and tradeoffs in the current implementation of `SL_specdecomp`. First, unlike `specparam`, `SL_specdecomp` does not currently include an iterative peak-finding algorithm that automatically proposes candidate rhythms and refines them. Instead, the user specifies candidate frequency bands for rhythms to be fit, and can compare candidate models using CVLL. This design has an important advantage: it encourages explicit hypothesis specification and can prevent overfitting by constraining rhythms to plausible bands. However, it is also a practical limitation in exploratory settings, where automated peak detection is useful.

Second, our simulations were mostly designed to test model misspecification in parameter regimes where the true decomposition is identifiable. This was intentional: if the broadband and rhythmic parameters are intrinsically weakly identifiable, then poor recovery may reflect the structure of the inverse problem rather than the specific probability or true-spectrum model being tested. Result 3 is an exception, where we show that `specparam` but not `SL_specdecomp` confounds rhythms and broadband when the broadband component below the knee is weakly identifiable, a scenario that is likely to occur in real data. We leave a more systematic study of weakly identifiable or non-identifiable regimes—for example, overlapping rhythms, very low-amplitude peaks, broad peaks that resemble the aperiodic component, or short noisy windows—to future work.

Third, our likelihood treats the frequency bins as approximately independent Gamma observations (after sub-sampling). This approximation is appropriate in the regime targeted here: sufficiently long time windows, multitaper smoothing, and frequency subsampling at approximately the multitaper resolution, where spectral leakage and finite-window correlations between frequencies are expected to be small. However, the model does not explicitly account for spectral leakage or for the correlations between nearby frequency bins induced by estimating spectra from finite time series. This approximation will break down for short time windows, strongly nonstationary signals, or spectra with sharp features whose energy spreads across neighboring frequencies. A natural extension would be to replace the conditionally independent Gamma likelihood with a likelihood or covariance model that explicitly accounts for finite-window spectral leakage and frequency–frequency correlations. We leave this extension to future work.

Overall, we provide evidence that two pervasive model misspecifications — a multiplicative true spectrum model and Gaussian, homoscedastic residual assumptions — can systematically bias rhythmic and broadband decomposition of neural power spectra. Correcting the probability model to reflect Gamma-distributed, heteroscedastic sampling variability substantially improves parameter recovery and predictive accuracy, and using an additive true spectrum model aligns the decomposition with standard biophysical superposition models of field potentials. These results imply that widely interpreted broadband findings, such as anesthesia-associated slope changes, can be attenuated or otherwise distorted by rhythm– broadband confounding under misspecified models. `SL_specdecomp` offers a principled alternative that respects both the physical generative intuition and the statistical structure of spectral estimators, providing a more reliable foundation for biomarker-oriented spectral analysis.

## Acknowledgments

We thank Becky Belisle, Thomas Donoghue, Yongho Lim, Will Liberti, and Eric Denovellis for valuable feedback on this work.

## Data and Code Availability

The empirical ECoG data analyzed in this study are publicly available through Neurotycho.org and were originally described by Yanagawa et al. (Yanagawa et al., 2013) and Nagasaka et al. (Nagasaka et al., 2011). The spectral decomposition framework introduced here is available as the open-source package `SL_specdecomp` at https://github.com/Stephen-Lab-BU/SL_specdecomp. Gaussian-process simulations with analytically specified power spectra are implemented in `SL_GPsim`, available at https://github.com/Stephen-Lab-BU/SL_GPsim. Code used to reproduce the manuscript analyses and figures is available at https://github.com/Stephen-Lab-BU/Bloniasz_Stephen_Estimation.

## Author Contributions

Patrick F. Bloniasz and Emily P. Stephen contributed equally in all parts of the Conceptualization, Methodology, Software, Formal analysis, Funding acquisition, Data curation, Visualization, Writing, Review, and Editing.

## Supplementary Section 1: Simulating Gaussian processes from a specified true spectrum

To benchmark spectral decomposition methods, we require simulated time series for which the ground-truth power spectrum is known. In our study, the ground truth is defined directly in the frequency domain by a target spectrum *S*_*µ*_(*f* ), rather than by assuming a particular time-domain model such as an autoregressive or Ornstein–Uhlenbeck process. This is important because it allows us to generate signals with arbitrary broadband and rhythmic structure, making the simulations agnostic to any one particular theory of what real neural power spectra should look like. We can then estimate a power spectrum from the simulated signal and ask whether a spectral decomposition method correctly recovers the known generating components.

Given a target two-sided spectrum *S*_*µ*_(*f* ), we simulate a mean-zero stationary Gaussian process using the approximate frequency-domain method of (D.Percival, 1992). This construction is useful here because it can be applied to a wide class of continuous spectral shapes while remaining computationally simple. The key idea is to build Fourier coefficients whose variances match the desired power at each frequency, impose the conjugate symmetry required for a real-valued signal, and then transform these coefficients back to the time domain.

Let *S*_*µ*_(*f* ) denote the desired two-sided spectrum of a real-valued, mean-zero stationary Gaussian process. For a real-valued signal, the spectrum is symmetric in frequency. We evaluate *S*_*µ*_(*f* ) on a dense uniform grid

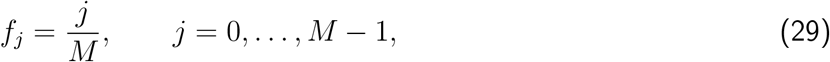

where *M* is even and chosen much larger than the desired sample length *N* . In our simulations, we set *S*_*µ*_(0) = 0, corresponding to a mean-zero process. We then generate independent standard normal random variables *W*_*j*_ and construct complex Fourier coefficients

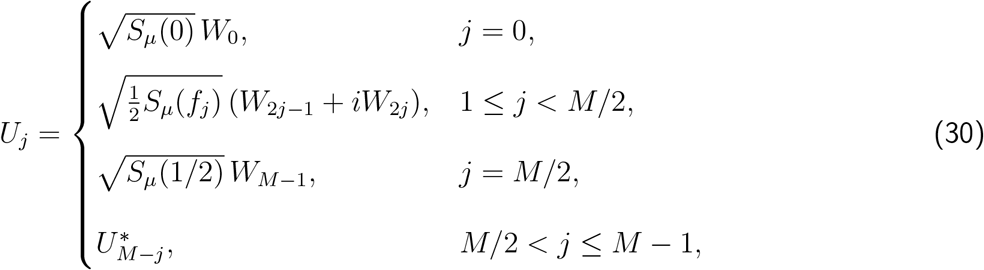

where 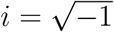 and ∗ denotes complex conjugation. The conjugate symmetry in Eq. (30) ensures that the inverse transform is real-valued.

The simulated time series is then obtained by an inverse discrete Fourier transform:

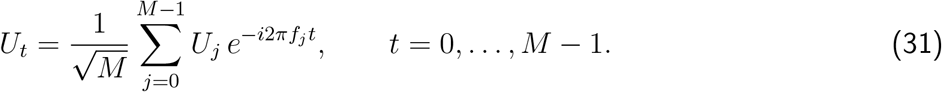

By construction, {*U*_*t*_} is a real-valued Gaussian process. Its autocovariance is determined by the target spectrum evaluated on the discrete frequency grid:

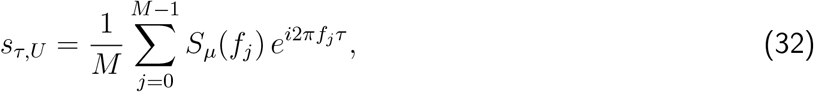

which is the discrete-frequency analogue of the usual inverse Fourier relationship between covariance and spectrum,

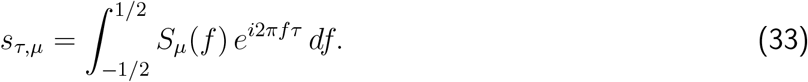

The practical point is that the frequency grid has to be sufficiently fine in order for the autocovariance of the simulated process to closely match the autocovariance implied by *S*_*µ*_(*f* ). If the frequency grid isn’t fine enough (e.g. if it has the same number of points as the desired timeseries, *M* = *N*, a common choice), then the periodicity implied by the inverse discrete Fourier transform leads to undesired covariances between time points (see (D. Percival, 1992) for more detail). So we choose *M* ≫ *N* to achieve a fine frequency grid, where in our implementation *M* corresponds to `n_fft` and *N* to `n_samples`. If `n_fft` is not provided manually, we set it to the next power of 2 greater than or equal to max(*f*_*s*_*/*Δ*f, N* ), where Δ*f* is the target frequency resolution. We then retain only the first *N* time-domain samples.

This simulation strategy is well suited to the goals of this paper. First, it allows us to generate signals from arbitrary ground-truth spectra, so our benchmarks do not depend on assuming that neural spectra arise from a particular parametric time-domain process. Second, because the simulated process is Gaussian with known true spectrum, the resulting direct spectral estimates have the Gamma sampling properties derived in the Background section. Third, defining the ground truth directly in the frequency domain makes it possible to test spectral decomposition methods in the same domain in which they operate, and therefore to evaluate directly whether they correctly separate broadband and rhythmic structure.

## Supplementary Section 2: Spectral decomposition model in SL_specdecomp

This appendix summarizes the probabilistic models implemented in our open-source package `SL_specdecomp`. The goal of the package is to decompose the spectrum into broadband and narrowband (rhythmic) components. We outline both the *additive* and *multiplicative* variants, specifying the functional forms of the components, the priors on their parameters, and the probability distribution for the observations (i.e. the likelihood).

### Broadband and rhythmic predictors

For a set of frequencies *f* up to the Nyquist frequency, we denote the underlying broadband spectrum by *P*_BB_(*f* ) and the contribution from *j* rhythmic peaks by *P*_rh,*j*_(*f* ). In `SL_specdecomp` these functions are constructed from two predictors:

1. **Lorentzian (“1/f”)** Each broadband component is modeled as a Lorentzian (“1/*f* “-like) function

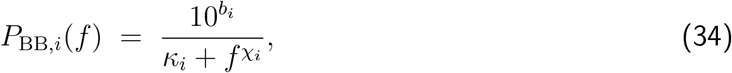

where *b*_*i*_ sets the offset in log-10 power, *χ*_*i*_ is the spectral exponent (the negative slope on a log–log plot), and *κ*_*i*_ encodes a knee frequency such that the spectrum flattens below the knee. Multiple broadband components can be summed to capture a spectrum with more than one knee.
2. **Mirrored Gaussian** Each rhythmic bump is represented by a mirrored Gaussian window

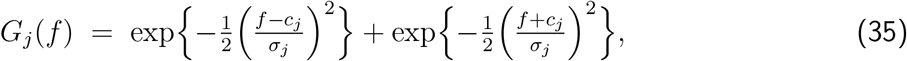

where *c*_*j*_ is the center frequency of the *j*th rhythm and *σ*_*j*_ controls its spread. The mirrored form enforces symmetry about zero and reduces edge effects at the Nyquist frequency. A scaling factor specific to each band is used in the additive model to put the rhythmic amplitudes on a comparable scale.

### Additive power model

The additive model reflects the physical superposition of independent generators and writes the latent power spectrum as a sum of the broadband background and narrowband bumps:

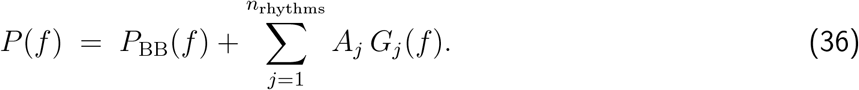

Here 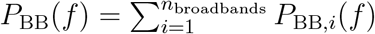 is a sum of Lorentzians from Eq. (34) and *A*_*j*_ is the linear amplitude of the *j*th Gaussian bump.

#### Default priors

The package uses weakly informative default priors that encode qualitative expectations about spectral shape while remaining permissive. For each broadband component:

- The slope parameter *χ*_*i*_ has a truncated normal prior on slope_*i*_ = −*χ*_*i*_ with mean −2 and standard deviation 1, truncated to (−5, −0.5). This centers the slope around −2 (consistent with many empirical spectra) while allowing steeper or flatter values.
- The knee frequency *f*_knee,*i*_ has a logarithmic uniform prior between 1 Hz and the Nyquist frequency. The parameter 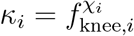 in Eq. (34) is then deterministically computed from *χ*_*i*_ and *f*_knee,*i*_.
- The offset *b*_*i*_ has a normal prior with mean equal to the median of log_10_-power in the data and standard deviation 5. This centers the background level near the observed spectrum.

For each rhythmic bump in the additive model:

- The center *c*_*j*_ is uniformly distributed within a user-specified band (or within canonical bands such as delta, theta, etc.).
- The bandwidth *σ*_*j*_ follows a truncated normal prior with mean 3 Hz, standard deviation 2 Hz, and bounds (0.5, 12) Hz.
- The linear amplitude *A*_*j*_ is positive. By default, we compute a band-specific reference scale *s*_*j*_ as the 50th percentile of the observed linear power within rhythm band *j*, and place a normal prior on log_10_ *A*_*j*_ centered at log_10_ *s*_*j*_ with standard deviation 1.25. This scale is used only to center the prior; the observed spectrum itself is not rescaled.

### Multiplicative power model

In some cases users may prefer to model narrowband structure as multiplicatively modulating the background, analogous to existing log-space methods. The multiplicative model in `SL_specdecomp` writes

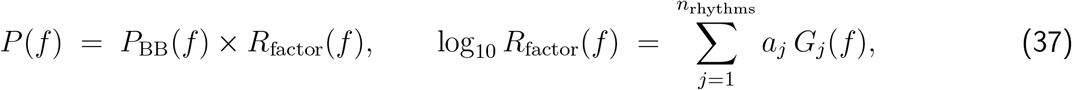

with the same Lorentzian form for *P*_BB_(*f* ). The quantity *a*_*j*_ now controls the height of the *j*th bump in log-power units.

#### Priors

For the broadband parameters the priors are as above, except that the offset *b* has a wider normal prior (*σ* = 10) to accommodate multiplicative scaling. For each rhythmic bump:

- The center *c*_*j*_ and bandwidth *σ*_*j*_ have the same priors as in the additive model.
- The log–amplitude *a*_*j*_ has a half-normal prior with scale parameter 1.

### Likelihood model and inference

Let 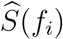 denote an averaged direct spectral estimate formed from *K* uncorrelated direct estimates. As reviewed in the main text, these estimates are approximately *Gamma* distributed. We therefore model each observed datum 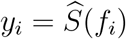 as

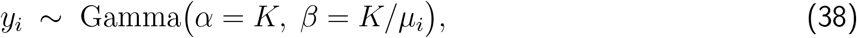

where *µ*_*i*_ = *P* (*f*_*i*_) is the latent mean power at frequency *f*_*i*_ under either the additive model (36) or the multiplicative model (37). Here we use the shape–rate parameterisation of the Gamma distribution: *α* is the shape parameter and *β* is the rate parameter, so the corresponding scale parameter is 1*/β* = *µ*_*i*_*/K*. Under this convention, 𝔼 [*y*_*i*_] = *α/β* = *µ*_*i*_ and 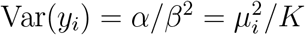, matching the heteroscedasticity of averaged direct spectral estimators. Here *K* is the number of uncorrelated direct estimates being averaged: for spectra averaged only across independent tapers, *K* = *n*_tapers_; for Welch spectra, *K* is the number of uncorrelated segments; for trial-averaged spectra, *K* is the number of independent trial spectra. More generally, *K* may be treated as an effective number of uncorrelated direct estimates.

#### Fitting the models

The models described above are expressed in the probabilistic programming language PyMC. Each model is fully specified by the choice of the number of broadband and rhythmic components, their allowed frequency bands, and whether sparse rhythms are used. Given observed frequency–power pairs (*f*_*k*_, *y*_*k*_), the posterior distribution over all parameters is sampled with the No-U-Turn Sampler (NUTS) by default; more optimized solutions are also supported, such as BlackJAX. Posterior summaries such as posterior means, credible intervals, and posterior predictive checks can then be computed. Users may customise priors on any parameter by passing specification dictionaries to the modelling functions.

## Supplementary Section 3: Figure-generation methods and model settings

### General spectral-estimation settings

Unless otherwise noted, spectra were estimated from signals sampled at 1000 Hz using multitaper spectral estimation on non-overlapping 30 s windows with time–half-bandwidth product *NW* = 2 and 3 tapers. For analyses of broadband biomarkers, the slope was computed by ordinary least-squares regression on the parametric fit of the power spectrum between 40–60 Hz, log_10_(power) on log_10_(*f* ). High-*γ* power was summarized as mean power over 80–180 Hz in the parametric fit of the power spectrum. When cross-validated log-likelihood (CVLL) was evaluated, each 30 s window was partitioned into five non-overlapping 6 s folds, and held-out spectra were estimated with *NW* = 1 and 1 taper. In the CVLL analyses, the training spectrum for each fold was averaged over the four non-held-out folds, so model fitting to the training spectrum used *K*_train_ = 4. The held-out spectrum was computed from one fold with one taper, so the predictive likelihood for the held-out spectrum used *K*_test_ = 1.

For peak-frequency summaries, we defined the recovered peak frequency as the continuous frequency within the relevant rhythm band at which the derivative of the fitted rhythmic curve is zero. This derivative-zero definition was used for the peak-frequency recovery panels in the simulation figures.

### Model-fitting settings

For `specparam`, all fits used `aperiodic_mode=‘knee’` and peak-width limits of 1–30 Hz. The minimum peak height was 0 a.u., which sets an absolute limit on the minimum height (above aperiodic) for any extracted peak. The peak threshold was 2 a.u.; “peak threshold” is the relative threshold above which a peak height must cross to be included in the model. The maximum number of allowed peaks was matched to the intended number of rhythms in each benchmark wherever possible; for example, if there are two ground truth rhythms, `specparam` was set to look for at most 2 rhythms, but because the user cannot enforce how many peaks to fit it can fit fewer. For `SL_specdecomp`, fits used one broadband component and a figure-specific number of rhythmic components. Posterior inference was performed with BlackJAX, or the user can use NUTS (no-u-turn sampling) using two chains and target acceptance probability 0.90. Draw and tuning counts were reduced only for cross-validation refits.

For comparison in Sup. Fig. 1, we also computed a naive broadband slope estimate that did not rely on spectral decomposition. Specifically, for each 30 s non-overlapping window, we estimated the multitaper power spectrum using the same settings as in Fig. 7 (*NW* = 2, 3 tapers), restricted the analysis to the positive-frequency grid used for the Figure 6 comparisons, and then fit an ordinary least squares line to log_10_ power as a function of log_10_ frequency over 40–60 Hz. The slope of this line was taken as the naive slope estimate.

**Supplementary Table 1.** Common spectral-estimation settings used across figures unless otherwise noted.

| Setting | Value |
| --- | --- |
| Sampling rate | 1000 Hz |
| Default window length | 30 s |
| Default multitaper parameters | $NW = 2, 3$ tapers |
| Primary analysis band | 1–200 Hz for simulation benchmarks; 0.1–200 Hz for awake/anesthesia decompositions |
| Broadband slope band | 40–60 Hz |
| High-γ band | 80–180 Hz |
| Cross-validation folds | 5 folds of 6 s each |
| Cross-validation multitaper parameters | $NW = 1, 1$ taper |

### Monkey ECoG dataset

For the analyses in Fig. 1, Fig. 7, Sup. Fig. 1, and Sup. Fig. 2 we used publicly available electrocorticography (ECoG) recordings from the *Macaca fuscata* monkey dataset (M1) hosted on Neurotycho.org. The experimental recording protocol is described in (Yanagawa et al., 2013), and the surgical implantation procedures and electrode coverage are described in (Nagasaka et al., 2011). Chronic ECoG electrodes were placed over the left hemisphere, with coverage spanning frontal, parietal, temporal, and occipital cortical regions.

**Supplementary Table 2.** Figure-specific data sources and simulation settings. “—” indicates that the figure used empirical data rather than a standalone ground-truth simulation. Figure 1 is a composite overview and therefore references the source analyses for its empirical and simulation panels.

| Figure | Duration | Exponent | Offset | Knee (Hz) | Data source / rhythmic component specification |
| --- | --- | --- | --- | --- | --- |
| 1 | Mixed | Mixed | Mixed | Mixed | Composite overview. Empirical panels use macaque ECoG during propofol anesthesia. Simulation panels illustrate the Gamma sampling distribution and a simulated-anesthesia decomposition example; final imported decomposition panels are sourced from the corresponding Figure 7 empirical payload and Figure 8 simulation payload rather than a single standalone Figure 1 simulation. |
| 2 | 60 s | 2.0 | 0.5 | 90 | Broadband-only benchmark. |
| 3 | 30 s | 2.0 | 0.5 | 60 | Additive ground-truth decomposition benchmark. One rhythm centered at 8 Hz with amplitude 2 and Gaussian width parameter $\sigma = 3$ Hz; evaluated across repeated simulations. |
| 4 | 30 s | 2.0 | 0.5 | 60 | Multiplicative ground-truth decomposition benchmark using the same broadband and rhythm settings as Figure 3, but with multiplicative/specparam-like true spectrum structure. One rhythm centered at 8 Hz with amplitude 2 and Gaussian width parameter $\sigma = 3$ Hz. |
| 5 | 30 s | 2.0 | 0.5 | 60 | Rhythm-height sensitivity benchmark using the Figure 3 additive ground truth. One rhythm centered at 8 Hz with Gaussian width parameter $\sigma = 3$ Hz; height varied over $\{0.5, 1.0, 2.0, 4.0\}$. |
| 6 | 30 s | 2.0 | 0.5 | 60 | Cross-validated log-likelihood benchmark using the same Figure 3–4 simulation regimes. CVLL compares specparam, Gamma-multiplicative SL_specdecomp, and Gamma-additive SL_specdecomp under additive and multiplicative ground truth. |
| 7 | — | — | — | — | Empirical macaque ECoG during eyes-closed wakefulness and propofol anesthesia. Representative 30 s windows and all 30 s windows from eyes-closed wakefulness ($n = 28$, after excluding two artifact windows from 30 original windows) and propofol anesthesia ($n = 18$, no windows excluded) are decomposed with specparam and additive SL_specdecomp. |
| 8 (awake) | 900 s | 2.533 | 5.240 | 57.420 | Ground-truth simulation generated from the posterior-mean Figure 7 awake SL_specdecomp fit. Two rhythms: 16.047 Hz ($A_{lin} = 421.040$, $\sigma = 3.161$ Hz) and 26.214 Hz ($A_{lin} = 24.978$, $\sigma = 5.642$ Hz). Synthetic windows are independent 30 s windows generated from identical ground-truth parameters, with 30 independent 30 s synthetic windows. |
| 8 (anesthesia) | 540 s | 3.508 | 6.994 | 19642.776 | Ground-truth simulation generated from the posterior-mean Figure 7 anesthesia SL_specdecomp fit. Three rhythms: 0.453 Hz ($A_{lin} = 41398.482$, $\sigma = 1.342$ Hz), 16.741 Hz ($A_{lin} = 134.815$, $\sigma = 3.270$ Hz), and 25.661 Hz ($A_{lin} = 42.726$, $\sigma = 6.032$ Hz). Synthetic windows are independent 30 s windows generated from identical ground-truth parameters, with 18 independent 30 s synthetic windows. |
| Sup. 1 | — | — | — | — | Empirical macaque ECoG; full-session 40–60 Hz slope trajectories for naive OLS, specparam, and additive SL_specdecomp, with exact Figure 7 target windows and aligned state-specific slope summaries. |
| Sup. 2 | — | — | — | — | Empirical macaque ECoG; window-wise CVLL comparison between matched additive SL_specdecomp models with 2 rhythms (without 20–30 Hz) versus 3 rhythms (with 20–30 Hz). |

**Supplementary Table 3.**
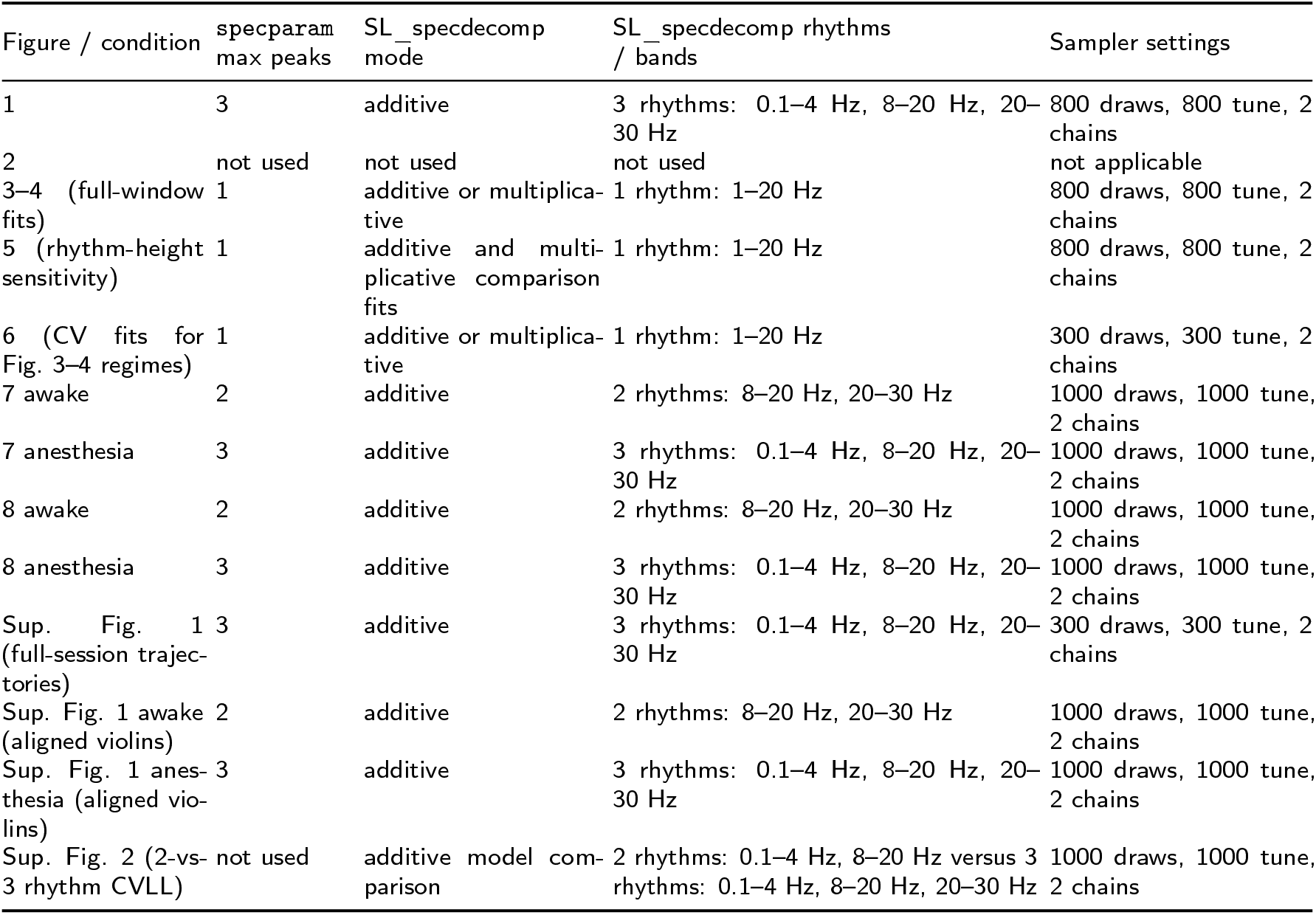
Decomposition settings for specparam and SL_specdecomp.

Recordings were acquired at a sampling rate of 1 kHz. In the session analyzed here, neural activity was first recorded for 20 min during the awake condition before propofol administration. Propofol was then infused at 5.2 mg kg^−1^ until loss of behavioral consciousness (LOC). LOC was operationally defined as the absence of a behavioral response to hand manipulation or to tactile stimulation of the nostril or philtrum with a cotton swab (Yanagawa et al., 2013). After LOC was established, ECoG was recorded for an additional 10 min while propofol administration was maintained, after which the infusion was stopped and the animal was allowed to recover.

For the empirical ECoG analyses in Fig. 1, Fig. 7, Sup. Fig. 1, and Sup. Fig. 2, we performed a visual outlier-inspection pass on the corresponding raw traces and multitaper spectra. This inspection identified two visually obvious artifact windows in the awake condition: original window indices 7 and 4, corresponding to 3.25–3.75 min and 1.75–2.25 min relative to the start of the analyzed awake-eyes-closed segment. These two awake windows were excluded from the state-specific decompositions, representative-fit selection, all-window overlays, peak rasters, broadband summary metrics, and CVLL analyses unless explicitly marked for visualization. No anesthesia windows were excluded. Thus, the state-specific analyses used 28 awake 30 s windows and 18 anesthetized 30 s windows.

### Additional robustness analyses for the real ECoG comparison in Fig. 7

Sup. Fig. 2 reports an additional model-comparison analysis for the real macaque ECoG data, using the same visual-outlier exclusions as the main state-specific decompositions. We compared two matched additive `SL_specdecomp` models within each state: a 2-rhythm model without a 20–30 Hz rhythm band, and a 3-rhythm model that includes the 20–30 Hz rhythm. Predictive performance was quantified by window-wise cross-validated log-likelihood (CVLL), using the same five-fold cross-validation scheme described above. Together with Sup. Fig. 1, this indicates that explicitly modeling the 20–30 Hz rhythm improves predictive fit during anesthesia.

Sup. Fig. 1 complements Fig. 7 by placing the slope analyses back into the full recording session. The top panel shows full-session 40–60 Hz slope trajectories computed on 30 s non-overlapping windows for naive OLS, `specparam`, and additive `SL_specdecomp`, with dashed lines marking the event labels and gray shading denoting the exact awake-eyes-closed and anesthetized intervals used for the state-specific analyses. The bottom panels show the exact state-specific slope distributions used for the Fig. 1 and Fig. 7 comparisons. Square markers denote the representative target windows used for the main-text Fig. 7 panel fits. These full-session and aligned state-specific summaries show that the anesthesia-associated steepening of the 40–60 Hz slope remains visible in the full-session context and in the state-specific summaries, and that this state shift is larger for additive `SL_specdecomp` and naive OLS than for `specparam`.

---

**Supplementary Figure 1.**
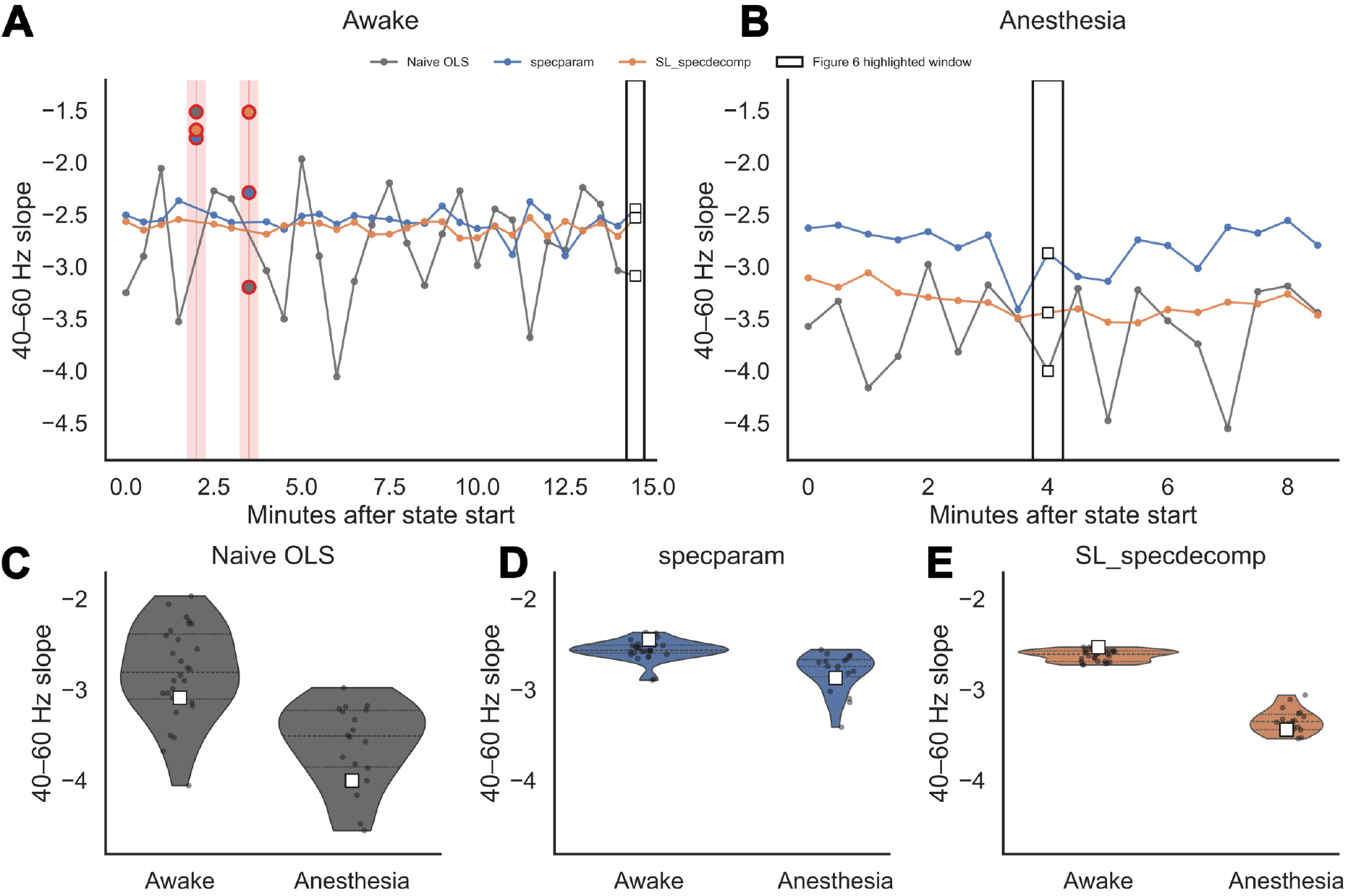
State-specific slope trajectories show the Fig. 7 analysis windows and preserve the anesthesia-related slope differences. (A,B) Window-wise 40–60 Hz slope trajectories for macaque ECoG during the awake-eyes-closed state (A) and propofol-anesthesia state (B), computed from 30 s non-overlapping windows. Each trajectory compares a naive log–log ordinary least-squares estimate applied directly to the multitaper spectrum, the aperiodic slope inferred by specparam, and the broadband slope inferred by additive SL_specdecomp. Red shaded intervals and red-outlined points in (A) mark the two visually identified awake artifact windows that were excluded from the window analyses; no anesthesia windows were excluded. Black outlined rectangles and square markers denote the exact representative target windows used for the Fig. 7 fits. (C–E) State-specific window slope distributions for naive OLS (C), specparam (D), and additive SL_specdecomp (E), with individual windows overlaid as points and the Fig. 7 representative windows marked by square markers. The 40–60 Hz slope is more negative in anesthesia than in wakefulness, and this anesthesia-associated steepening is larger for naive OLS and additive SL_specdecomp than for specparam.

**Supplementary Figure 2.**
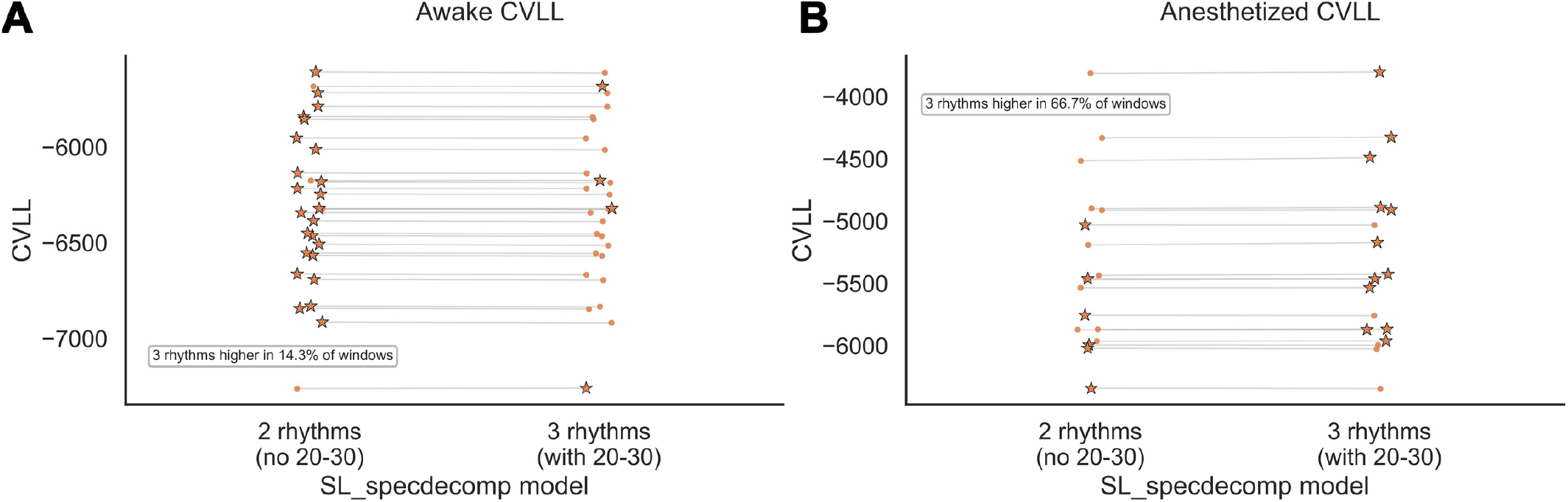
Inclusion of a 20–30 Hz rhythm improves additive SL_specdecomp fit in the anesthetized but not awake state. (A,B) Window-wise cross-validated log-likelihood (CVLL) comparison for macaque ECoG during awake eyes-closed rest (A) and propofol anesthesia (B), using two matched additive SL_specdecomp models fit to the same 30 s windows. The left column in each panel is a 2-rhythm model without the 20–30 Hz rhythm; the right column is a 3-rhythm model with the 20–30 Hz rhythm included. Each connected pair denotes one window, and star markers denote the higher-CVLL model for that window. The 3-rhythm model attains higher CVLL in 14.3% of awake windows and 66.7% of anesthetized windows, indicating that inclusion of the 20–30 Hz rhythm improves predictive fit primarily during anesthesia.

